# The evolution of a counter-defense mechanism in a virus constrains its host range

**DOI:** 10.1101/2022.04.14.488369

**Authors:** Sriram Srikant, Chantal K. Guegler, Michael T. Laub

## Abstract

Bacteria use diverse immunity mechanisms to defend themselves against their viral predators, bacteriophages. In turn, phages can acquire counter-defense systems, but it remains unclear how such mechanisms arise and what factors constrain viral evolution. Here, we experimentally evolved T4 phage to overcome a phage-defensive toxin-antitoxin system, *toxIN*, in *E. coli*. Through recombination, T4 rapidly acquires segmental amplifications of a previously uncharacterized gene, now named *tifA*, encoding an inhibitor of the toxin, ToxN. These amplifications subsequently drive large deletions elsewhere in T4’s genome to maintain a genome size compatible with capsid packaging. The deleted regions include accessory genes that help T4 overcome defense systems in alternative hosts. Thus, our results reveal a trade-off in viral evolution; the emergence of one counter-defense mechanism can lead to loss of other such mechanisms, thereby constraining host range. We propose that the accessory genomes of viruses reflect the integrated evolutionary history of the hosts they infected.

**Highlights:**

- Experimentally evolved T4 to overcome *E. coli toxIN*, a phage-defensive TA system
- Discovered TifA, a phage-encoded protein inhibitor of ToxN
- Amplification of the *tifA* locus drives large deletions elsewhere in the T4 genome
- Deleted genes in evolved T4 clones include those necessary to infect alternative hosts

## Introduction

Bacteria face the frequent threat of phage predation and have consequently evolved a diverse arsenal of anti-phage defense systems (Bernheim and Sorek, 2020). These defense systems have, in turn, driven the selection of counter-defense mechanisms in phage, underscoring the intense arms race between phages and their bacterial hosts (Samson et al., 2013; Hampton et al., 2020). There are several well-characterized examples of anti-restriction modification (RM) and anti-CRISPR proteins (Stanley and Maxwell, 2018). However, there is an ever-growing inventory of phage defense mechanisms in bacteria and how phages counteract these diverse defense systems is poorly understood (Samson et al., 2013). More generally, it remains unclear how phages can rapidly acquire resistance when confronted with a new defense system and what types of mutations and mechanisms are responsible. Probing how phage overcome bacterial defense systems will help reveal the molecular basis of bacteria-phage coevolution and may inform efforts to develop phage therapies for treating antibiotic-resistant bacterial infections.

An emerging class of potent phage defense genes are toxin-antitoxin (TA) systems, which typically comprise a protein toxin that is directly neutralized by its cognate antitoxin (Harms et al., 2018). For defensive TA systems, phage infection activates or triggers the release of the toxin, which can then block phage development (Guegler and Laub, 2021). Although the initially infected cell typically does not survive, particularly if the toxin also inhibits host cell processes during infection, these defensive TA systems prevent the release of new, mature virions, thereby preventing spread of an infection through a population.

How can a phage evolve to overcome a defensive TA system? There are three general possibilities: (i) a phage prevents activation of the TA system, (ii) a phage becomes resistant to the action of the toxin, or (iii) a phage acquires or modifies a factor that can inhibit or degrade the toxin. As TA systems may be activated by and target core phage processes, the latter is a potentially powerful mechanism because it does not require mutations in essential phage genes. Notably, like eukaryotic viruses, phages often harbor many ‘accessory genes’ that are not formally required for phage propagation in a naïve host in permissive conditions. For instance, T4 encodes nearly 300 proteins, only 62 of which are required to produce a new virion in common laboratory conditions (Miller et al., 2003). The remaining genes are mostly of unknown function but their presence in many related T4-like phage suggests they may play key roles in host specificity by overcoming host defense systems or enabling replication in specific growth conditions.

There are several examples of phage-encoded anti-TA factors. The T4 gene *dmd* encodes a direct inhibitor of the RnlA toxin from the *E. coli* K12 TA system RnlAB (Otsuka and Yonesaki, 2012; Wan et al., 2016). Consequently, RnlAB only protects *E. coli* against T4 phage lacking *dmd* (gene *61.5*). More recently, T-even phages including T4 were shown to encode an inhibitor (AdfA, encoded by gene *61.2* in T4) of DarT toxins (LeRoux et al., 2021). In *Pectobacterium atrosepticum*, the type III TA system *toxIN,* involving an endoribonuclease toxin and RNA antitoxin, protects against infection by several phages, including <ΔTE and <ΔA2. Selection for <ΔTE mutants that overcome the *P. atrosepticum toxIN* system identified phages that either expressed additional copies of a locus resembling the *toxI* antitoxin or that had acquired *toxI* from the *toxIN* system directly (Blower et al., 2012). This study indicates that phages may be poised to readily evolve mechanisms to overcome defensive TA systems.

Here, we evolved T4 phage to overcome a *toxIN* homolog from a natural plasmid in an environmental *E. coli* isolate that can provide *E. coli* MG1655 with potent defense (Fineran et al., 2009; Guegler and Laub, 2021). Using an experimental evolution approach adapted from that used in early phage therapy initiatives (Appelmans, 1921; Burrowes et al., 2019), we find that T4 can rapidly overcome *toxIN* through the segmental amplification and increased expression of *tifA* (previously 61.4), which encodes a protein inhibitor of ToxN. Strikingly, these segmental amplifications led to T4 genome instability, likely because an increased genome size is incompatible with the fixed capsid size of T4. The evolved phages subsequently acquired compensatory deletions that restored genome size, with the set of deleted genes varying among replicate evolved populations. These deletions compromise the ability of T4 to infect other strains of *E. coli,* often because the lost genes encode factors that overcome other, strain-specific defense systems. In sum, our work reveals the genome dynamics underlying the emergence of an anti-defense mechanism in T4 and demonstrates an evolutionary trade-off in which selection to infect one host can compromise infection of others.

## Results

### T4 can evolve to overcome *toxIN-*mediated defense in *E. coli*

We previously identified and characterized *toxIN_Ec_* (hereafter *toxIN* for simplicity), a type III TA system from an environmental isolate of *E. coli* that strongly protects *E. coli* MG1655 against T4 infection (Figure 1A; Guegler and Laub, 2021). To evolve T4 to overcome *toxIN*, we adapted the Appelmans protocol originally developed in phage therapy work to evolve phage able to replicate on resistant pathogenic hosts (Appelmans, 1921; Burrowes et al., 2019; Mapes et al., 2016). Briefly, our approach involved replicating T4 on both a sensitive host (-*toxIN*), which maintains the phage population size and generates diversity, and on a resistant host (+*toxIN*) to select for phages that can overcome the defense system (Figure 1B). During each round of evolution, serial dilutions of the phage population (six serial 10-fold dilutions producing a range of 10^7^ to 10^1^ phage) were each inoculated with 10^5^ host cells to produce a range of multiplicities-of-infection (MOIs, defined as the ratio of phages to bacteria) from 10^2^ to 10^-4^ in 96-well plates and then grown for 16-20 hrs. This protocol allows phage to evolve by a combination of point mutations and recombination events within a phage genome, as well as recombination between co-infecting phage. The clearing of cultures across MOIs provided a visual readout of the phage population evolving to infect the resistant host (Figure 1B, S1A). Throughout the evolution protocol, the T4 population titer remained at 10^5^-10^6^ pfu/µL, with an estimated 1-3 phage infection generations occurring in wells from high to low MOI within a single round of evolution (see Methods). We set up 5 replicate evolutions of T4 to overcome *toxIN* carried on a medium-copy plasmid in *E. coli* MG1655 (+*toxIN*), and a control population replicating only on -*toxIN* cells (MG1655 with an empty vector) (Figure S1A). We plated serial dilutions of the phage populations after each round on lawns of +*toxIN* cells to identify *toxIN-*resistant phage clones as they arose and fixed (Figure S1B).

**Figure 1.**
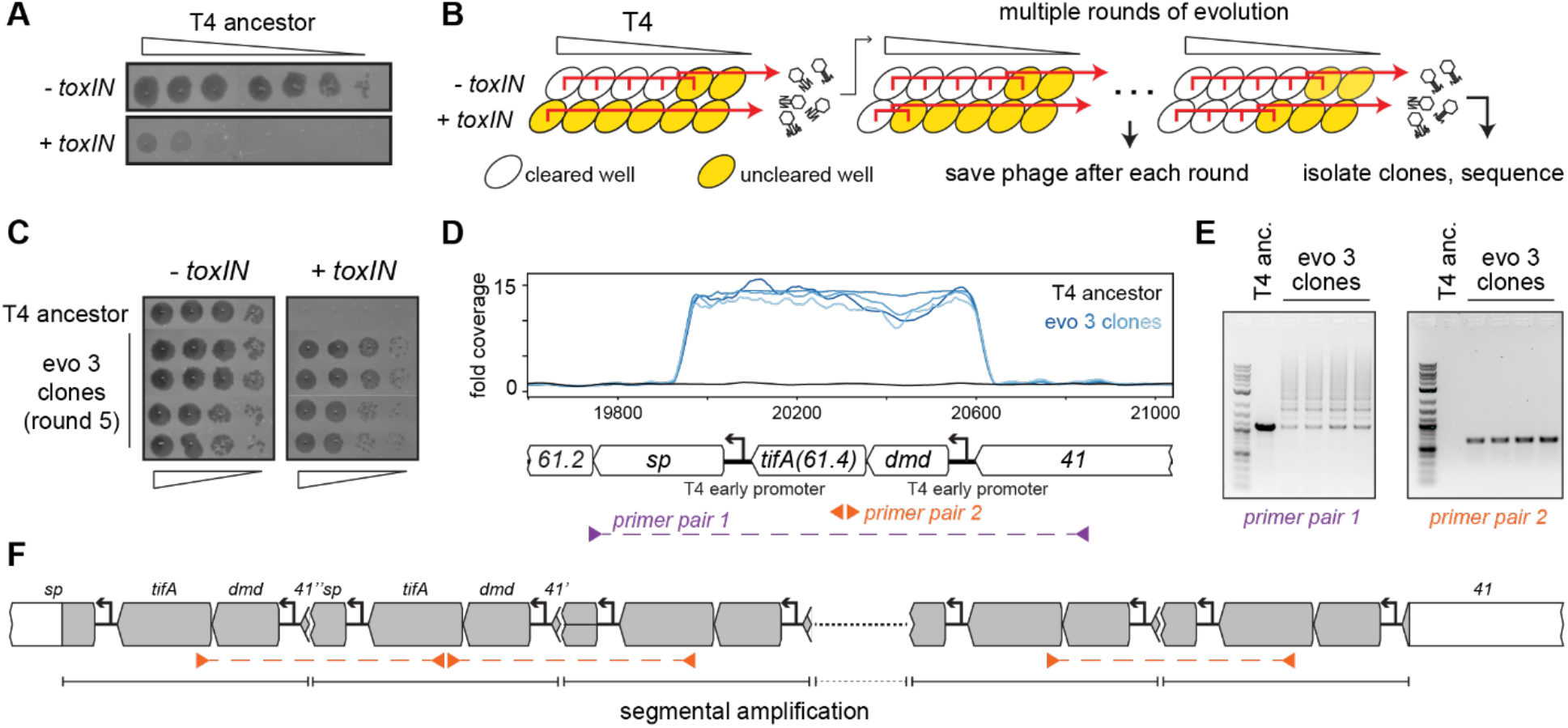
T4 rapidly evolves to overcome *toxIN* by segmental amplification of the *dmd-tifA(61.4)* locus. (**A**) T4 is restricted by +*toxIN E. coli*. Serial dilution plaquing assay of T4 spotted on -*toxIN* and +*toxIN* cells. (**B**) Phage evolution protocol to select for T4 that overcomes *toxIN*-mediated defense. Serial dilutions of T4 are used to infect *+toxIN* and *-toxIN* cells at multiple MOIs. The ancestral T4 only clears wells containing *-toxIN* cells. Phage from all cleared wells, and the last uncleared well, are pooled and used to inoculate the next round of evolution. As the population evolves, it can also infect and clear *+toxIN* cells. Phage populations from each round are saved to maintain a fossil record; individual clones are isolated on *+toxIN* cells and sequenced to identify mutations leading to *toxIN* resistance. (**C**) Comparison of plaquing efficiency for ancestral T4 and T4 isolates from one of the evolved populations after 5 rounds on *-toxIN* and *+toxIN* cells. (**D**) Read coverage at the *dmd-tifA(61.4)* locus following genome sequencing of multiple clones from one of the evolved populations. Primer pairs used in panel E to interrogate the segmental amplification are shown below the locus map. (**E**) PCR products using the primer pairs indicated in panel D for ancestral T4 and evolved clones. (**F**) Schematic of segmental amplification of *dmd-tifA* locus in evo 3 clones highlighting binding of primer pair 2 producing a cross-repeat product.

For one of the populations (evo 3), phage that could form plaques on lawns of +*toxIN* cells arose after 5 rounds of evolution, an estimated 5-15 infection generations. From this population, we isolated phages from four individual plaques on +*toxIN* lawns and confirmed that these evolved phages plaque as efficiently on +*toxIN* cells as on the control strain *-toxIN* (Figure 1C). To identify the mutations responsible for overcoming *toxIN*, we fully sequenced the genomes of these four evolved clones. No individual gene was mutated in all of the evolved clones. However, our sequencing revealed increased read coverage of a two-gene locus in the genome of each evolved clone (Figure 1D). This locus consists of two genes*, dmd (61.5)* and *61.4*, driven by an early T4 promoter (Liebig and Rüger, 1989; Miller et al., 2003). As noted above, Dmd is an inhibitor of the RnlA toxin of the RnlAB TA system (Otsuka and Yonesaki, 2012) and *61.4* encodes a putative 85 amino-acid protein of unknown function, which we have renamed *tifA* (<u>T</u>oxN inhibitory factor <u>A</u>) based on the studies below.

The increased read coverage of the *dmd-tifA* locus could indicate the presence of multiple copies of this locus scattered throughout the T4 genome or a local, segmental amplification of this locus. PCR using primers flanking this locus produced a ladder of products, indicating a segmental amplification generating tandem repeats (Figure 1E-F, S1D). PCR using divergent primers in the middle of the amplified region generated a band for the evolved clones but not the T4 ancestor (Figure 1E). This band is produced by primers annealing to neighboring repeats and thus reflects the size of a single repeat in the amplification.

### TifA is a protein inhibitor of ToxN

We hypothesized that the segmental amplifications in our T4 escape mutants led to increased expression of either *dmd* or *tifA*, allowing the phage to overcome *toxIN*. To determine which gene was responsible, we cloned *dmd* and *tifA* separately into a vector containing an inducible promoter and transformed each plasmid into +*toxIN* cells. We then infected both strains with the ancestral, wild-type T4 in the presence of inducer and compared their efficiency of plaquing (EOP) (Figure 2A). The ability of *toxIN* to inhibit T4 infection was completely lost following induction of *tifA*, suggesting that this gene was sufficient to prevent ToxN-mediated defense. Overexpressing *dmd* did not allow T4 to replicate in the presence of *toxIN*, indicating that this gene did not contribute to escape, but was likely included in the segmental amplification as the promoter for *tifA* lies upstream of *dmd* (Figure 1D).

**Figure 2.**
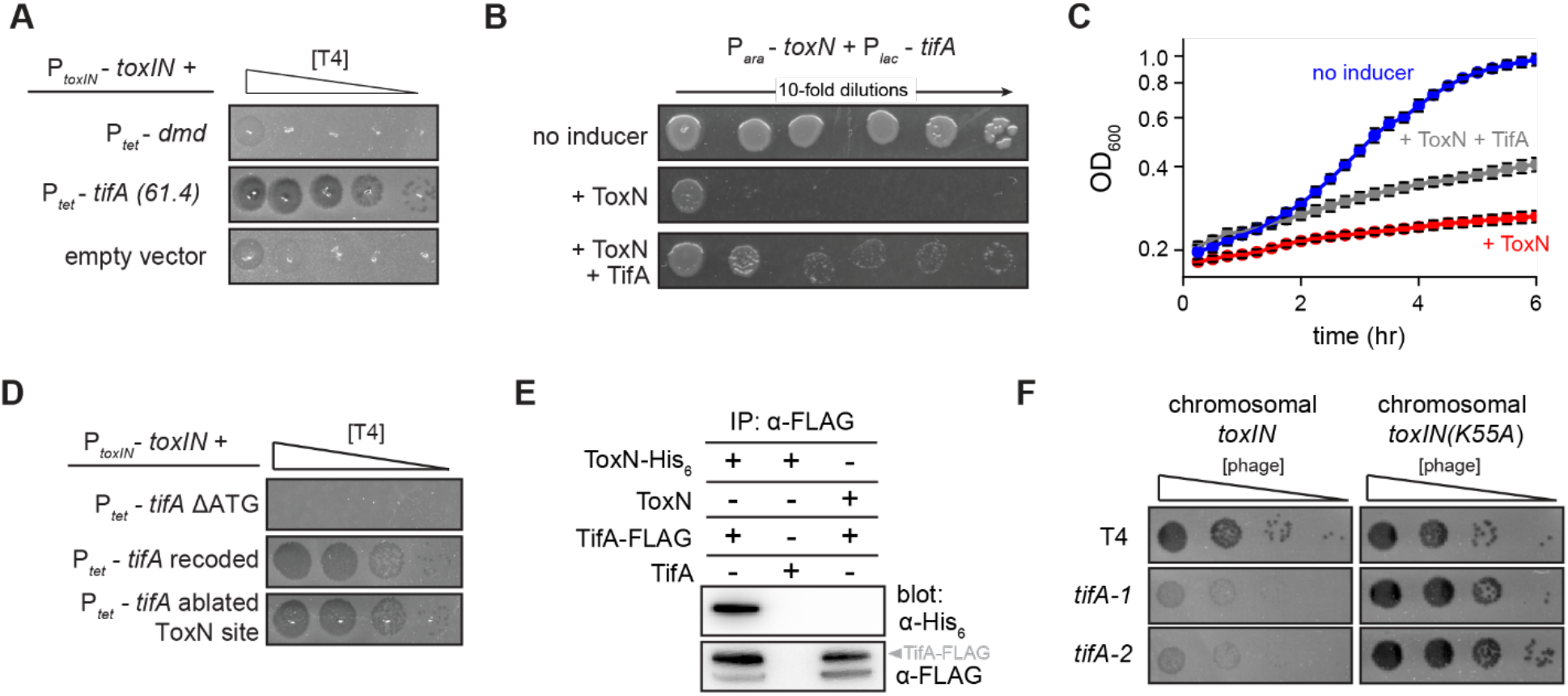
TifA (gp61.4) inhibits ToxN toxicity. (**A**) Serial dilution plaquing assay of T4 ancestor spotted on +*toxIN* cells also harboring a plasmid containing *dmd* or *tifA(61.4)*, or an empty plasmid. Anhydrotetracycline was added to the plates to induce *dmd* or *tifA*. (**B-C**) Representative plating assay (B) and growth curves (C) showing TifA rescue of ToxN. Plasmids harboring *toxN* and *tifA* under arabinose- and IPTG-inducible promoters, respectively, were transformed into *E. coli* MG1655. ToxN, ToxN and TifA, or neither were induced as indicated. (**D**) Serial dilution plaquing assay of T4 ancestor spotted on *+toxIN* cells containing plasmids expressing the *tifA* variants indicated. (**E**) Western blot of ToxN-His_6_ and TifA-FLAG (or untagged controls) following an anti-FLAG coimmunoprecipitation of T4-infected cells co-producing ToxN and TifA. The arrowhead highlights the band with a molecular weight matching that expected for TifA-FLAG. The identity of the band below is unknown but may represent a processed or truncated portion of TifA-FLAG. (**F**) Serial dilution plaquing assay of T4 ancestor and two *tifA* mutants on *E. coli* MG1655 with chromosomally-encoded *toxIN* or *toxIN*(K55A), which harbors an active-site mutation in ToxN.

Our results suggested that *tifA* is a phage-encoded factor that, upon overexpression, can offset ToxN activity during T4 infection. To test whether *tifA* encodes an antitoxin for ToxN, we cloned *toxN* into a vector that allowed for its inducible expression and co-transformed this plasmid into *E. coli* MG1655 with a plasmid containing *tifA* controlled by a separate, inducible promoter. As expected for a bacteriostatic toxin, inducing ToxN in uninfected cells inhibited the growth of *E. coli.* Co-induction of *tifA* was sufficient to partially rescue this toxicity, on both solid media and in culture (Figure 2B-C) indicating that *tifA* is a T4-encoded antitoxin for ToxN.

Given that the native antitoxin *toxI* is an untranslated RNA that binds and neutralizes ToxN, we sought to determine whether *tifA* encodes a protein or an RNA antitoxin of ToxN (Figure 2D). Expressing a version of *tifA* in which the start codon was deleted (*61.4* ΔATG) could no longer restore T4 infection of cells harboring *toxIN,* suggesting that *tifA* must be translated to function. Additionally, we found that a recoded variant of *tifA* that has a different mRNA sequence (55 of 258 nucleotides changed) but produces the same protein sequence could still restore T4 infection of +*toxIN* cells, indicating that the protein product of *tifA* was sufficient for ToxN inhibition. Finally, to verify that the *tifA* mRNA does not act as a competitive substrate for inhibition of ToxN, an endoribonuclease, we ablated a putative ToxN cleavage site at the 3′ end of *tifA*. Expression of this variant did not interfere with inhibition of ToxN, confirming that it is the protein product, and not the mRNA, of *tifA* that counters *toxIN* in our evolved T4 clones. It is these results that prompted us to rename gene *61.4* to *tifA* (<u>T</u>oxN inhibitory <u>f</u>actor <u>A</u>).

### TifA interacts with ToxN *in vivo*

To test whether TifA interacts directly with ToxN, we generated a strain producing ToxN from the *toxIN* locus with a C-terminal His_6_ tag and TifA with a C-terminal FLAG tag. We first verified that these tags did not affect activity of either protein (Figure S2A). We then infected cells with T4, lysed the cells, and performed an immunoprecipitation with anti-FLAG antibodies to isolate TifA and any interacting proteins. Western blotting with an anti-FLAG antibody confirmed that TifA was present, as expected, and blotting with an anti-His_6_ antibody demonstrated that ToxN was also recovered (Figure 2E). These results suggest that TifA interacts with ToxN, either directly or indirectly, *in vivo*.

If T4 encodes an inhibitor of ToxN, why is *toxIN* normally able to defend against wild-type T4? We reasoned that, because *toxIN* is encoded on a medium-copy (∼20 / cell) plasmid in our system, the single, native copy of *tifA* in T4 might not be sufficient to overcome ToxN. To test this idea, we introduced either *toxIN* or *toxIN(K55A),* which produces an enzymatically dead ToxN, into the *E. coli* MG1655 chromosome and infected both strains with T4 (Figure 2F). In this case, *toxIN* was no longer sufficient to protect against wild-type T4 infection. To confirm that this infection requires *tifA*, we used a CRISPR-Cas9 system to generate two mutant strains of T4 (see Methods) containing either a 98 bp deletion or 5 bp insertion disrupting the *tifA* open reading frame (Figure S2B). Both T4 mutant strains exhibited an ∼100-fold decrease in plaquing efficiency on a strain containing chromosomal *toxIN*, but were comparable to the wild-type T4 when tested on a *toxIN(K55A)* control strain (Figure 2F). These results suggest that native *tifA* expression is sufficient to overcome ToxN produced from the chromosome and that the segmental amplification that arose during our evolution experiments reflects a pressure to overcome the stronger expression of plasmid-borne *toxIN*.

### TifA homologs from other T4-like coliphage inhibit ToxN

Homologs of *tifA* are found adjacent to *dmd* homologs in the genomes of the coliphage T2, T6, and RB69, all of which are closely related to T4 (Figure 3A), and in other types of phages that infect a diverse set of bacterial hosts (Figure S3A). TifA homologs share a common architecture, with a central region containing a cluster of positively charged residues (Figure 3B, S3A). Although the T2, T4, and T6 homologs are nearly identical to one another, RB69 TifA is more divergent (56% identity) and lacks the C-terminal extension shared by the other three (Figure 3B). Interestingly, plasmid-borne *toxIN* strongly protects *E. coli* against T2, T4, and T6 infection, but offers substantially less protection against RB69, as measured by plaquing efficiency and cell death in shaking cultures (Figure 3B-C, S3B-C). Thus, we hypothesized that the more divergent TifA from RB69 may be a more potent inhibitor of ToxN.

**Figure 3.**
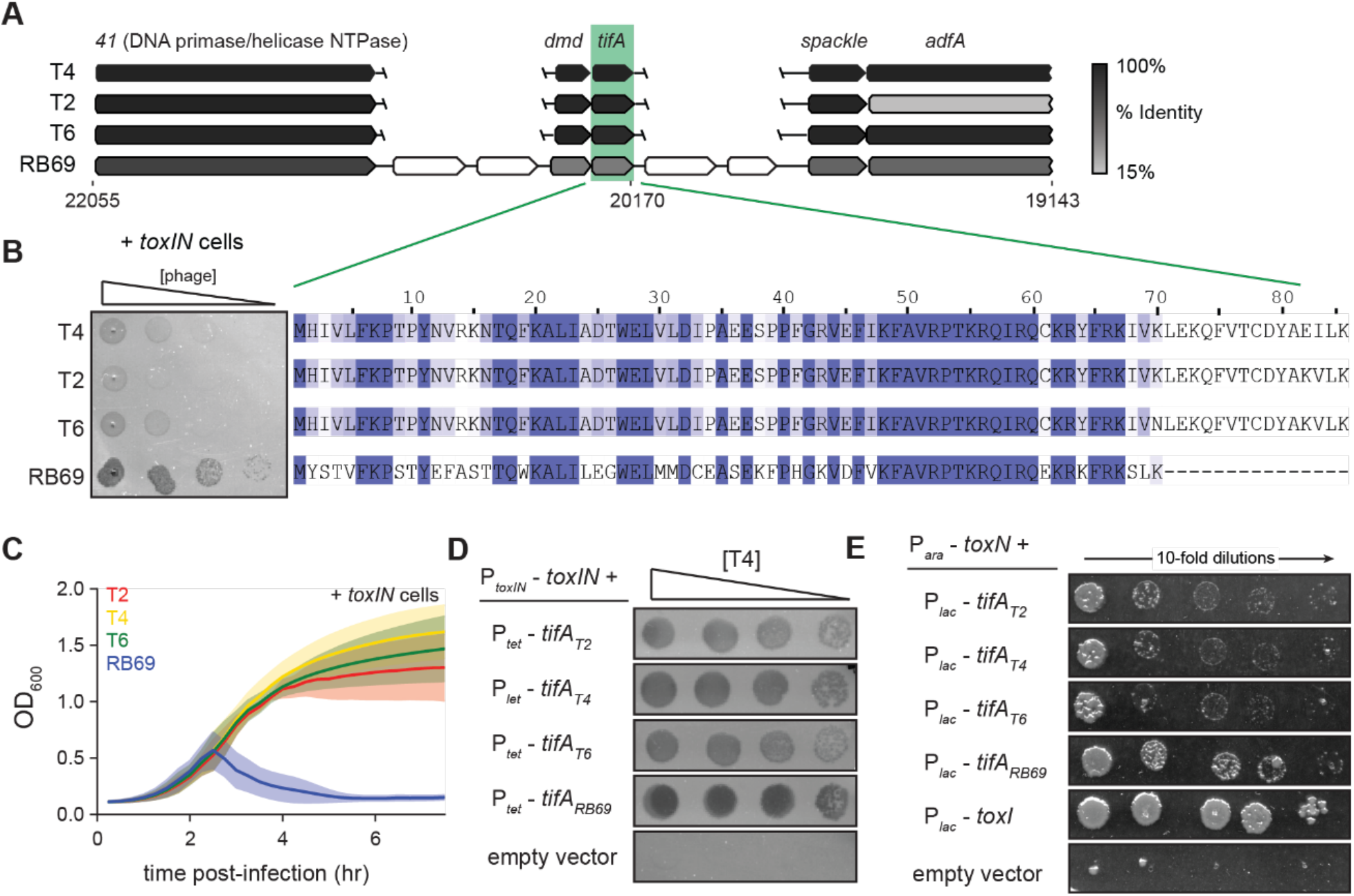
RB69 TifA is a more potent inhibitor of ToxN than the T4 homolog. (**A**) Schematic of syntenic region containing *tifA* in closely related T-even phages. Homologous genes are aligned vertically and colored by % identity to the homolog in T4. (**B**) (*Left*) Serial dilution plaquing assays of T4, T2, T6, RB69 on *+toxIN* cells. (*Right*) Multiple sequence alignment of TifA homologs from T4, T2, T6, and RB69. (**C**) Growth curves following infection of *+toxIN* cells with T4, T2, T6 or RB69 phage, each at a MOI of 10^-3^. (**D**) Serial dilutions of T4 ancestor spotted on *+toxIN* cells expressing *tifA* homologs from T2, T4, T6, RB69, or harboring an empty vector. (**E**) Serial dilutions of *E. coli* cells expressing *toxN* and the *tifA* homolog from T2, T4, T6, or RB69, *toxI*, or alone.

To test this hypothesis, we cloned each *tifA* homolog from T2, T4, T6, and RB69 into a plasmid with an inducible promoter and transformed those plasmids into +*toxIN* cells. We then compared the ability of wild-type T4 to infect each strain (Figure 3C). Overexpressing any of the four TifA variants allowed T4 to form plaques on +*toxIN* lawns, compared to the empty vector (Figure 3D), but with cells producing RB69 TifA producing the clearest plaques. We also tested each variant for protection against T2 infection, again finding that RB69 TifA produced the clearest plaques and led to the highest EOP on *+toxIN* lawns (Figure S3D). Finally, we expressed each *tifA* homolog in uninfected cells ectopically producing ToxN and found that all the TifA homologs rescued cells from ToxN-mediated toxicity to some extent (Figure 3E). We conclude that the TifA homolog from each of these T-even phages is a ToxN inhibitor, but that RB69 may encode a more potent inhibitor, enabling RB69 phage to be least affected by the *toxIN* system we tested.

### T4 genome evolution proceeds through frequent recombination events

For the phages from population 3 that were initially isolated and sequenced after 5 rounds of evolution, we noticed that individual plaques exhibited substantial variability in size (Figure S1C), suggestive of some genomic variation during isolation. We therefore continued the evolution for this and the other 4 replicate populations for 20 additional rounds, maintaining selection for infection of *+toxIN* cells. By plating serial dilutions of the evolving populations on *+toxIN* lawns, we could track, as they arose, T4 clones that could infect *+toxIN* cells in each population (Figure 4A-C, S4A). ToxIN-resistant phages emerged at a different round in each population, but with comparable levels of resistance stabilizing after 25 rounds (Figure 4A-B, S4A). We then sequenced the genome of one phage clone isolated from each population (Figure 4C). There were no point mutations in common across all 5 evolved phage clones. However, as with the first clones sequenced (Figure 1D), each of these sequenced genomes had an amplification of the *dmd-tifA* locus, as manifested by a ∼10-fold increase in read counts in this region (Figure 4D, S4B). For the clone from population 5, *dmd* is included in the amplification but not the promoter upstream. However, in this case there is a promoter downstream of *tifA* that ends up oriented in a manner that would allow it to drive expression of the *dmd-tifA* cistron following amplification (Figure S4C).

**Figure 4.**
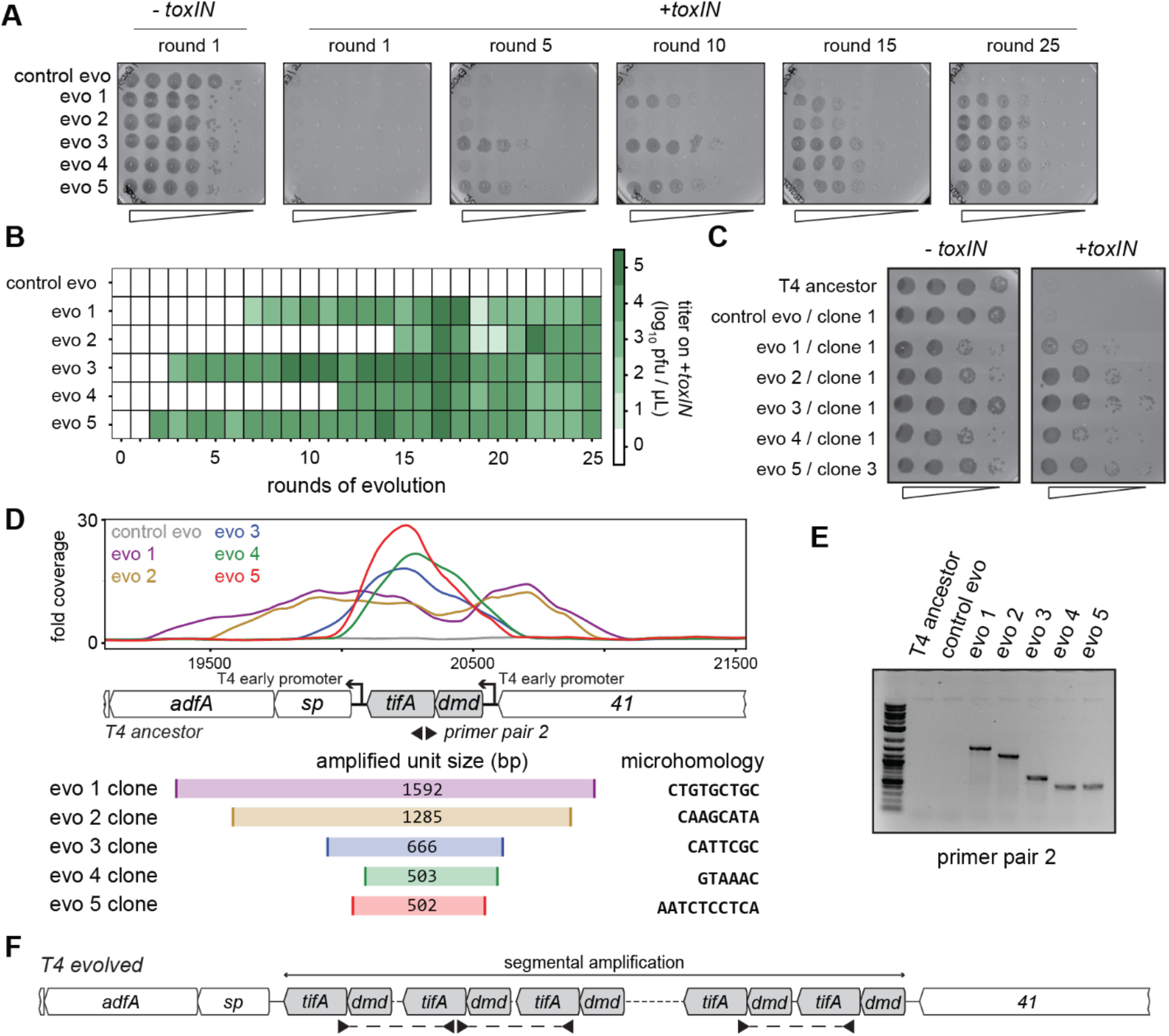
Unique recombination events generate *dmd-tifA* amplifications in replicate T4 evolution experiments. (**A**) Serial dilutions of evolving populations (1 control evolution and 5 replicate populations) after rounds 1, 5, 10, 15, and 25. (**B**) Heat map summarizing the titer of *toxIN-*resistant T4 as they arise in the independent evolving populations. Titer was estimated by serial-dilutions of evolving populations spotted on *+toxIN* cells (as in panel A). (**C**) Serial dilutions of isolated clones from evolved populations spotted on *-toxIN* and *+toxIN* cells. (**D**) Fold coverage from genome sequencing of evolved T4 isolates around *dmd* and *tifA*. (**E**) PCR with divergent primers, indicated in panel D, used to map the size of the repeated unit in the tandem segmental amplifications. (**F**) Schematic of segmental amplification of *dmd-tifA* locus in evo clones highlighting binding of primer pair 2 producing a cross-repeat product that depends on the specific location of the microhomology.

The precise boundaries of the segmental amplification differed in each evolved population. This finding indicates that the same adaptive solution, the amplification of the *dmd-tifA* locus, occurred independently in each evolved population. We performed a PCR with divergent primers that would only amplify a product following a segmental amplification to confirm the size of the amplified region in each population (Figure 4D-F). The different sizes of the band in clones from different populations confirmed that the repeat region was unique in clones of each population (Figure S4B). Sanger sequencing of the PCR products revealed that the boundaries of each segmental amplification coincided with unique, short (6-10 bp) regions of microhomology that normally flank the *dmd*-*tifA* locus (Figure 4D, S4D). In the evolved clones only one of the two regions of microhomology remain between neighboring repeats (Figure S4D). Thus, the segmental amplifications observed likely arose through homologous recombination between these regions of microhomology, as also seen previously in T4 (Kumagai et al., 1993; Mosig, 1987; Wu et al., 1991, 2021).

Strikingly, in addition to an amplification of the *dmd-tifA* locus, each evolved clone of T4 also had large deletions elsewhere in its genome (Figure 5A-B). Individual deletions were different in each clone and ranged from 0.7 to 6 kb in length, for a total of 5 to 11 kb lost in each genome (Figure 5C). We confirmed the absence of each deletion by PCR using primers that flank the deleted region inferred by genome sequencing (Figure S5A). These deletions likely enable T4 to maintain a constant genome size by acquiring deletions that compensate for the amplifications selected during evolution. The T4 genome is packaged by a headful mechanism in which 172 kb from a genomic concatemer is inserted into a pro-head. The inserted DNA represents 102% of the T4 genome (168 kb), with ∼3.3 kb of terminal redundancy (Grossi et al., 1983; Kim and Davidson, 1974; Rao and Black, 2005). To ensure stable packaging and inheritance of the genome, the segmental amplifications that arose following selection on +*toxIN* cells likely required the genome to contract elsewhere. Consistent with this interpretation, the size of the deleted region in each evolved clone was roughly proportional to the size of the segmental amplification, as estimated from the increased read coverage across the *dmd-tifA* region (Figure 5C).

**Figure 5.**
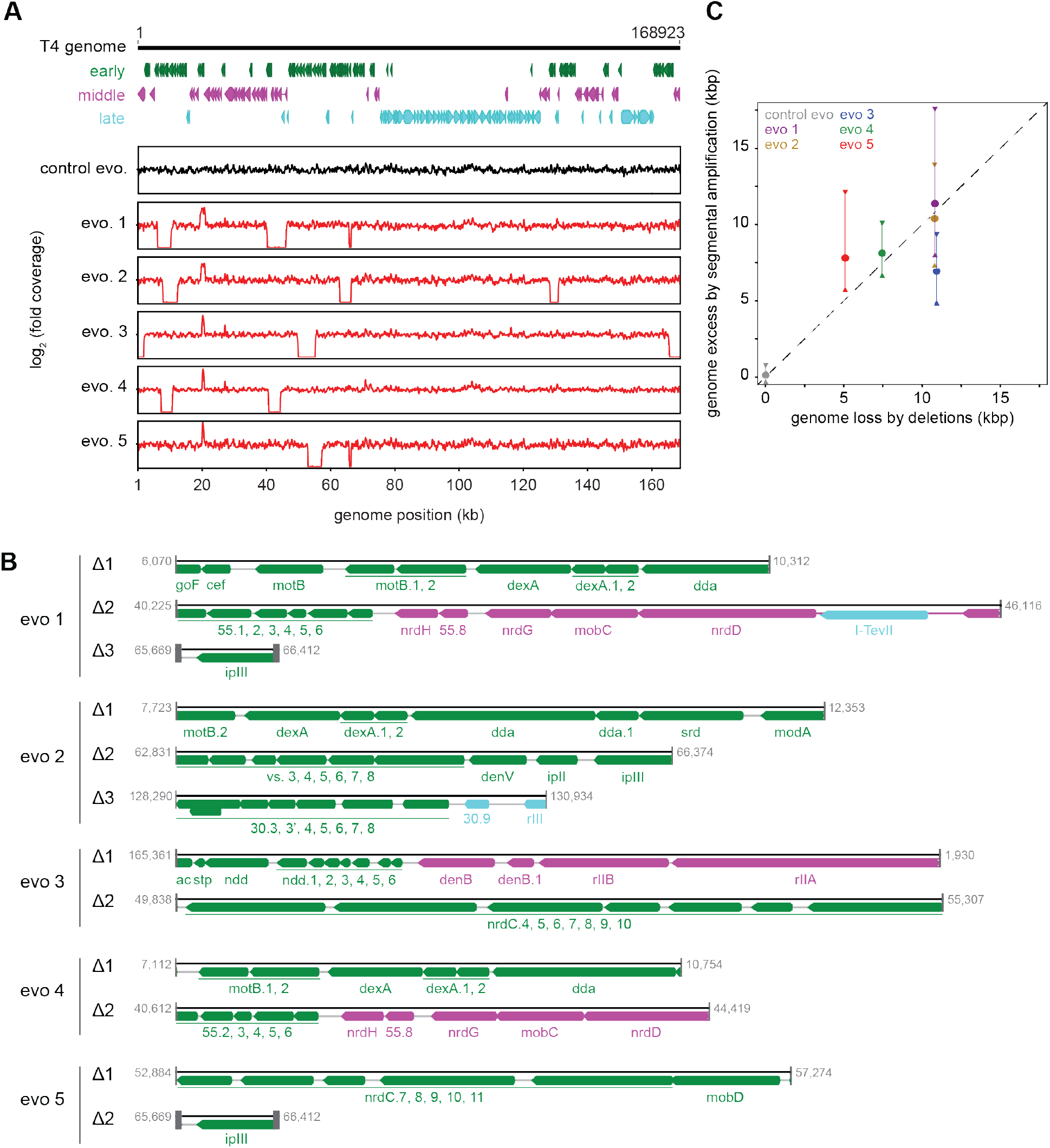
Segmental amplifications in the T4 genome are compensated by large genomic deletions. (**A**) Fold coverage calculated from genome sequencing of clones from replicate evolved populations plotted across the entire genome, showing regions of increased coverage (*dmd-tifA* locus) and large genomic deletions manifesting as a loss of coverage. Genes are colored based on their regulation during the phage life cycle; early (green), middle (magenta) and late (cyan). (**B**) Summary of deleted regions in each evolved clone show the flanking regions of microhomology (grey rectangles). (**C**) Scatter plot of increase in genome size estimated from fold-coverage across *dmd-tifA* repeat region (circle represents median and triangles represent interquartile range of coverage) against the size of genome deletion in clones from each population.

Our results indicate that T4 rapidly evolves through recombination between short regions of microhomology within its genome (Kumagai et al., 1993; Mosig, 1987; Wu et al., 2021). To overcome the *toxIN* system, such recombination events cause the segmental amplification of *dmd* and *tifA*. The resulting genome is then likely too big to be stably maintained, giving rise to the variability in genome coverage seen during intermediate rounds of our evolution, with different individual plaques containing different deletions that restore genome size (Figure S1C, S5B). At later rounds, one of these genome configurations rises to a high frequency in the population (Figure 5, S5B-C).

### Genome deletions prevent T4 from infecting other strains of *E. coli*

None of the 62 genes that are essential for producing new virions during infection of *E. coli* in rich media at 37 °C (Figure S6A) were included in the deletions identified in our evolved clones (Miller et al., 2003). However, we hypothesized that the genes lost in our evolved clones may compromise infection of other strains of *E. coli*.

One of the deletions that arose in evolved phage population 3 (evo 3) contained the *rII* locus (Figure 5B). The *rIIA* and *rIIB* genes allow T4 to replicate in *E. coli* strains harboring a λ-lysogen (Benzer, 1955). The RexAB system of λ allows it to exclude T4 *rII* mutants by a mechanism that remains poorly characterized (Wong et al., 2021). As predicted, the T4 clones that have deletions spanning *rII* lost the ability to replicate on a λ-lysogen of MG1655 (Figure 6A, S6B). Spot assays and whole-genome sequencing of clones isolated from different time-points during the evolution confirmed that this population of phage first acquired the ability to infect *+toxIN* cells and then subsequently lost the ability to replicate in a λ-lysogen once the genomic deletions including *rII* had fixed in the population (Figure S5B-C, S6B). Expressing *rIIA* and *rIIB* from plasmids in a MG1655 λ-lysogen partially restored the infectivity of an evolved T4 clone, confirming the causal nature of the *rII* deletion (Figure 6A). These results suggest that the genes lost to compensate for *dmd-tifA* amplification may include genes that help T4 overcome other defense and exclusion mechanisms, like RexAB.

**Figure 6.**
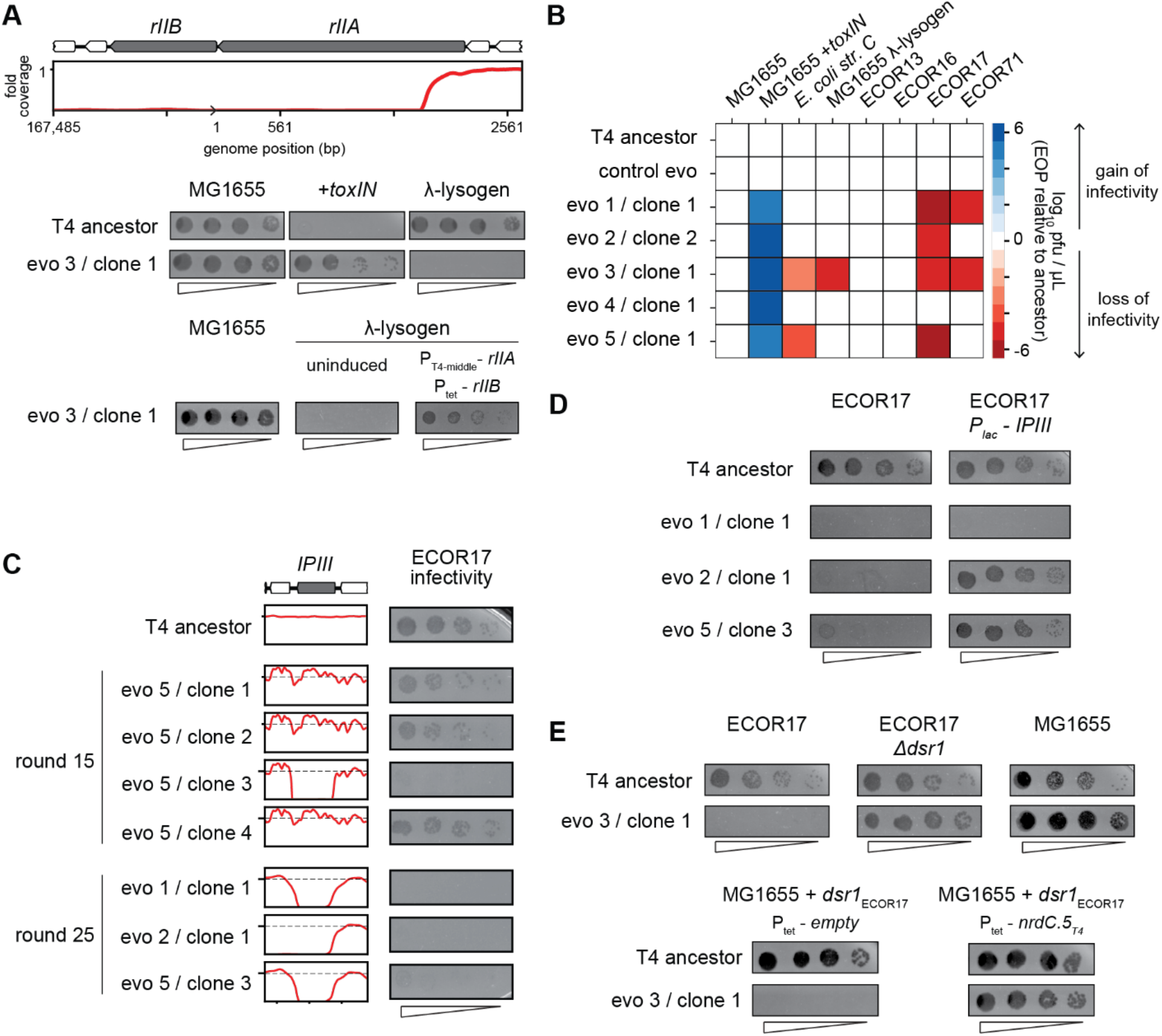
T4 genes lost during evolution are essential for replication in alternate hosts. (**A**) (*Top*) Fold coverage map for *rIIA* and *rIIB* locus in a clone from population 3 (evo 3). (*Bottom*) Serial dilutions of the T4 ancestor and evo 3 clone 1 on the strains indicated. (**B**) Heat map quantifying the relative efficiency of plaquing (EOP) of each evolved clone compared to the T4 ancestor for the indicated *E. coli* strains. (**C**) Fold coverage maps for the *IPIII* gene in the indicated clones of T4 from evolved populations evo 1, 2 and 5, with the corresponding serial dilutions of each clone on ECOR17. (**D**) Serial dilutions of evolved isolates with *IPIII* deletions on ECOR17 and ECOR17 + *IPIII* cells. (**E**) (*Top*) Serial dilutions of T4 ancestor and evo 3 clone 1 on ECOR17, ECOR17 11*dsr1*, and MG1655 lawns. (*Bottom*) Serial dilutions of T4 ancestor and evo 3 clone 1 on MG1655 with plasmid-borne *dsr1*_ECOR17_ also harboring a plasmid expressing *nrdC.5_T4_* or empty vector.

The *rII* deletions were only found in clones from one of the five evolved populations, and specifically affect infection of λ-lysogens. We therefore tested the evolved T4 clones for their ability to infect strains from the ECOR collection, a set of *E. coli* strains isolated from diverse environments that have similar core genomes but widely varying accessory genomes (Ochman and Selander, 1984). We focused on strains that can be infected by our ancestral T4 phage, which included 4 strains from this collection, ECOR13, ECOR16, ECOR17, and ECOR71, as well as *E. coli str. C* (Figure 6A, S6C). Clones from populations 1, 2, 3 and 5 (evo 1, 2, 3 and 5) failed to infect ECOR17 and clones from populations 1 and 3 (evo 1 and 3) failed to infect ECOR71 (Figure 6A). The loss-of-replication phenotype arose in each population after they had acquired the ability to overcome *toxIN*, indicating that deletions that compensate for the segmental amplifications led to the loss of genes required for infecting these hosts (Figure S6B, S6D).

To identify the gene(s) responsible for loss-of-replication, we asked which phage gene deletions correlated with the inability to infect ECOR17. The only gene lost in common across clones in evo 1, 2 and 5 was *ipIII*, implying that it enables T4 to infect ECOR17 (Figure 6C, 5B). T-even phages encode a set of proteins – including IPIII – called internal proteins: they contain a sequence motif that drives packaging into mature virions such that the proteins are injected into a host with the phage genome (Kutter et al., 1995; Leiman et al., 2003; Mullaney and Black, 1996). IPI inhibits the restriction-modification system *gmrSD* (Bair and Black, 2007), but the function of the other internal proteins is not known. We hypothesized that IPIII may also be an anti-defense protein that targets a host defense system. Indeed, providing *ipIII_T4_* on a plasmid in ECOR17 restored the ability of clones from evo 2 and 5 to infect this host (Figure 6D). We also found that having the host cells express a variant of IPIII lacking the capsid-targeting sequencing (*ipIII_T4_ΔCTS*) restored infectivity of clones from evo 2 and 5, indicating that IPIII functions in the cytoplasm of ECOR17 to enable T4 replication (Figure S6E). For the clone from evo 1, *ipIII* on a plasmid did not restore infectivity, suggesting that phages in this population lack an additional gene required to infect ECOR17.

To identify the relevant host defense systems in ECOR17, we used available annotation pipelines (Payne et al., 2021; Tesson et al., 2021) to predict phage-defensive operons in the ECOR17 genome, finding 6 potential systems, which we deleted, individually, from ECOR17 (Figure S6F). None of these deletions restored the infectivity of phage that lack IPIII, suggesting that an unknown defense system exists in ECOR17 and that IPIII enables T4 to overcome this system. Though evo 3 clone 1 also lost the ability to infect ECOR17, its deletions do not include IPIII. For this clone, a disruption of *dsr1* in ECOR17 restored infection (Figure 6E, S6F). Dsr1 contains a Sirtuin domain and likely blocks phage replication by disrupting cytoplasmic NAD+ levels (Garb et al., 2021). Expression of the ECOR17 *dsr1* gene from its native promoter on a medium-copy plasmid conferred protection to MG1655 against evo 3 clone 1 (Figure 6E). Thus, a T4 gene deleted in evo 3 clone 1 must overcome Dsr1 in the context of T4 ancestor infection. Expressing the uncharacterized T4 early gene *nrdC.5* under an inducible promoter restored infectivity of evo 3 clone 1 on MG1655 carrying *dsr1* (Figure 6E). Thus, the T4 genome contains multiple genes (including *ipIII* and *nrdC.5*) that enable the infection of multiple hosts by overcoming the suite of defense systems in these hosts. Our results support the notion that phage genomes are flexible and can rapidly amplify genetic material allowing them to overcome a specific defense system, but at the cost of losing genes that enable infection of other hosts (Figure 6A).

## Discussion

### TifA and an anti-TA island in T4-like phage

Bacteria and phages are locked in an ongoing coevolutionary battle, with extant genomes of each reflecting the outcomes of the underlying molecular arms race (Koskella and Brockhurst, 2014; Hampton et al., 2020). Given the fast generation times of bacteria and phages, experimental evolution offers an opportunity to study this arms race as it unfolds. Here, using an adapted version of Appelmans protocol (Appelmans, 1921; Burrowes et al., 2019; Mapes et al., 2016), we evolved T4 to overcome the type III TA system *toxIN*. Genome sequencing of evolved clones identified a two-gene operon *dmd-tifA* that was independently amplified in five replicate populations. We demonstrated that *tifA*, which encodes an 85 amino acid protein, inhibits ToxN, and that the amplification of *tifA* allows T4 to overcome plasmid-borne *toxIN* (Figure 7). Homologs of TifA are found in many T4-like phage, suggesting that these phage may frequently encounter hosts harboring *toxIN-*like systems. However, these homologs vary in both their sequence and their ability to inhibit ToxN, suggesting that each TifA may be ‘tuned’ to inhibit a particular, cognate ToxN of an ecologically relevant host.

**Figure 7.**
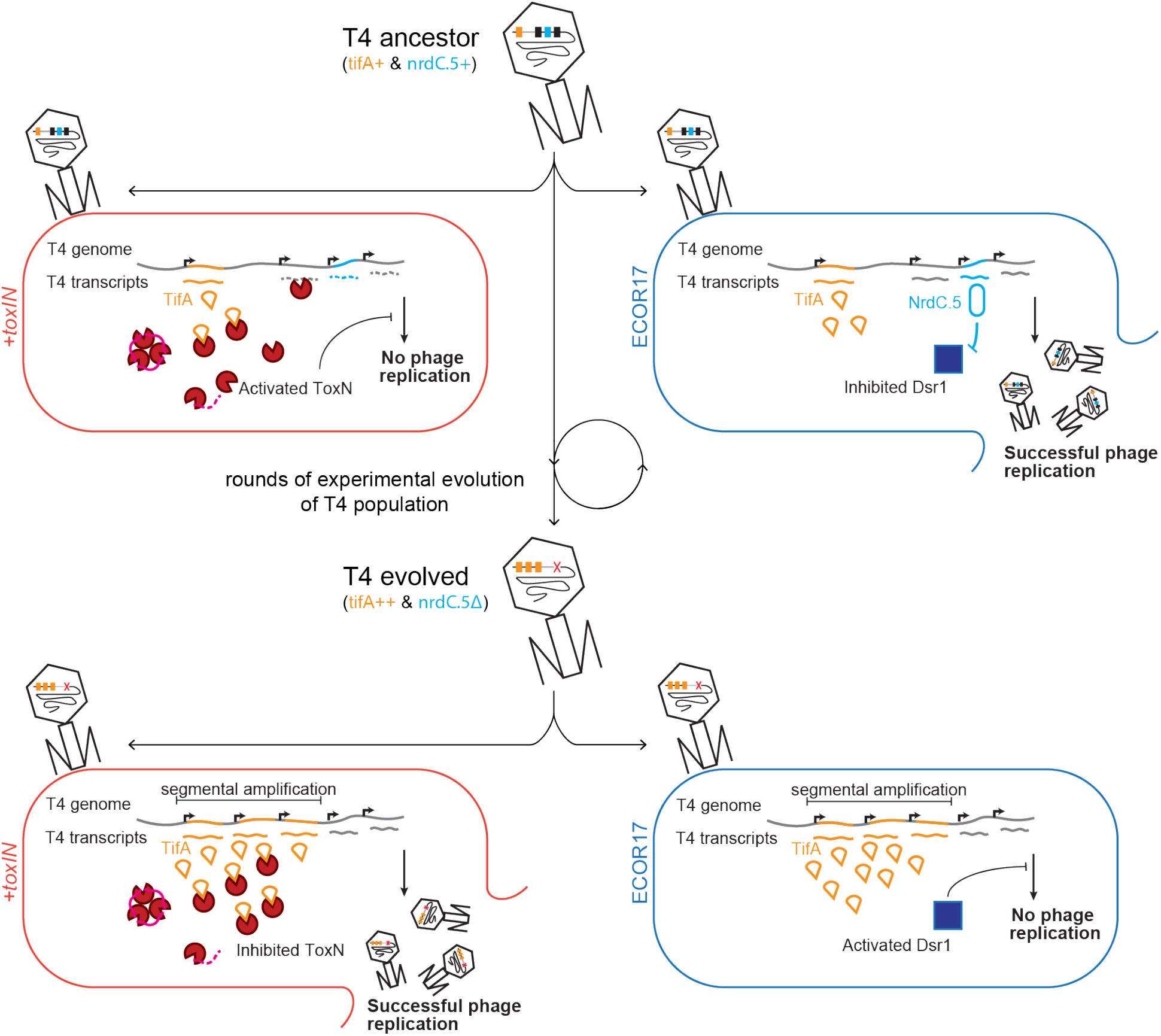
Model of T4 evolving to overcome toxIN by genomic variation affecting host range. The genome of T4 contains a number of accessory genes (*tifA*, *nrdC.5*, etc) that enable infection of hosts by overcoming hosts defenses (*toxIN*, *dsr1*, etc). T4 ancestor is unable to infect *+toxIN* cells due to inability to inhibit the action of ToxN but can infect an alternate host (ECOR17) using *nrdC.5* to overcome the bacterial defense gene *dsr1*. Experimental evolution to infect *+toxIN* cells produce T4 evolved clones that have segmental amplifications of the tifA-locus and compensatory deletions of non-essential genes like accessory genes needed to infect alternate hosts. A T4 evolved clone can thus infect *+toxIN* cells by having enough TifA to inhibit ToxN, but has lost *nrdC.5* in a deletion, and thus lost the ability to infect ECOR17.

TifA is analogous to the co-operonic gene *dmd*, which encodes a direct inhibitor of RnlA, the toxin of the *rnlAB* system in *E. coli*. Additionally, the nearby and conserved gene *61.2* (now *adfA*) was recently found to overcome the type IV TA system *darTG* (LeRoux et al., 2021). Thus, this region of the T4 genome may be an anti-TA island, similar to the anti-RM and anti-CRISPR islands found in other phages (Pinilla-Redondo et al., 2020). Other genes in this region of the T4 genome are attractive anti-toxin candidates.

### Recombination-mediated amplifications and deletions drive T4 genome evolution

T4 evolved to overcome *toxIN* by amplifying *dmd*-*tifA*, likely via recombination between short homologous sequences that flank this locus. These recombination events may arise during replication of the T4 genome or during the repair of DNA breaks. The unique amplifications that arose independently in our five replicates highlight the high rates of recombination-based mutagenesis in phage evolution. Similar recombination events between short (2-10 bp) regions of homology in T4 have been seen previously in studies of the *rII* locus (Kumagai et al., 1993), genes *16-18* (Wu et al., 1991) and escape from CRISPR-Cas defense (Wu et al., 2021).

Our work also indicates that T4 can undergo large genomic changes, through recombination between short regions of homology leading to loss of the intervening region (Figure 5A, S5A). Genome packaging in T4 proceeds via a headful mechanism (Rao and Black, 2005), so any substantial gain in DNA must be offset by comparably sized losses. Thus, the gains in genetic material resulting from amplification of *dmd-tifA* likely necessitated the loss of other, non-essential DNA to maintain a genome that could be fully packaged into the head. A similar pattern of gene amplification in the *16-18* region of T4 followed by deletion of the gene *alt* was previously reported (Wu et al., 1991).

Gene amplification is a major source of evolutionary innovation. For instance, antibiotic resistance in bacteria frequently arises through gene amplification. Seminal studies of *Salmonella* demonstrated that spontaneous tandem duplications arise frequently in bacterial genomes and can be present in as much as 3% of a population (Anderson and Roth, 1981, 1977). Gene duplications can provide intrinsic advantages, as with antibiotic resistance genes or in our case of *tifA*. Additionally, duplications create additional copies of a gene and hence provide a larger target for point mutations; if a beneficial mutation arises in any copy of the gene, it can provide a selective advantage and be maintained while other copies are lost by subsequent recombination. Such a scenario explains the apparently high rates of *lac*-reversions measured by Cairns and Foster when selecting for restoration of LacZ activity in strains harboring a point mutation in *lacZ* (Andersson et al., 1998; Cairns and Foster, 1991; Hendrickson et al., 2002). Similarly, a selection for poxviruses that can overcome the host restriction factor protein kinase R revealed an initial segmental amplification of the anti-host factor K3L, followed by subsequent acquisition of a beneficial point mutation in one copy K3L and subsequent loss of additional copies of the gene (Elde et al., 2012).

In the case of *tifA*, the entire T4 ancestor population likely contains a single-copy of the *dmd-tifA* locus, which is insufficient to overcome plasmid-borne *toxIN*. Thus, the segmental amplifications that increase *tifA* copy number could not have been identified through a direct, one-step selection for escape mutants. However, the recombination events leading to these amplifications arose within several rounds in each of our replicate populations, indicating that this is a readily available pathway to overcoming *toxIN*. We did not observe point mutations in *tifA* early on that improve its inhibition of ToxN, likely because segmental amplifications via recombination arise more frequently. We also did not observe beneficial point mutations arising after the segmental amplifications, leading to subsequent collapse of the amplification, as with protein kinase R. We speculate that the fitness gains and number of beneficial point mutations in *tifA* are likely too small to arise and fix.

### The evolution of phage host range

The genes deleted in our evolved T4 phage are all non-essential for infection of *E. coli* MG1655 grown in laboratory conditions, but may be crucial for T4 to infect other hosts. Indeed, we found that the deletions in four of our five replicate populations compromised infection of other strains of *E. coli*. The loss of *rII* led to an inability to infect Δ lysogens, consistent with prior work showing that *rII* is required to neutralize the λ-encoded anti-T4 system RexAB. Similarly, we found that IPIII, which was lost in three populations, is required for T4 to infect *E. coli* strain ECOR17. Internal proteins, including IPIII, are packaged with and then subsequently injected with the T4 genome. These proteins likely act against “first-response” defense systems like RM systems in each host. For instance, IPI is known to inhibit the *E. coli* type IV RM system *gmrSD*. Which defense system IPIII inhibits remains to be determined.

The accessory genomes of phages likely include a broad range of anti-defense genes that profoundly influence their host ranges. The set of anti-defense genes in a phage genome at a given time will reflect, in part, those that have recently been selected for based on the hosts that phage has infected. As our experimental evolution demonstrates, if selection to infect a host and overcome its defense systems is not maintained, the corresponding anti-defense genes, which are not essential for phage replication, can be lost following pressure to infect new and different hosts (Figure 7). Such genome dynamics, which stem from the relentless coevolution of phage with their hosts, may help to explain the staggering diversity of accessory genes within phage genomes (Pope et al., 2015) and their sometimes narrow host ranges across hosts with varied and dynamic immune profiles (Hussain et al., 2021; Kauffman et al., 2022).

### Concluding remarks

Our experimental evolution approach enabled the identification of new anti-phage defense and phage counter-defense mechanisms. In addition to finding TifA as a novel inhibitor of ToxN, the characterization of our evolved T4 isolates indicated that IPIII likely antagonizes a novel, as-yet unknown defense system in ECOR17. Additionally, we discovered a homolog of the recently identified Dsr1 system in ECOR17 that can be counteracted by the non-essential gene *nrdC.5* in wild-type T4. Collectively, these findings underscore how much remains to be discovered about the molecular arms race between bacteria and phages, which has profoundly shaped the genomes of both.

Phages evolve rapidly, but the forces and mechanisms that drive this evolution are poorly understood. Experimental evolution helps to reveal the tempo and mutations critical to phage genome evolution. Efforts to dissect how phages evolve to overcome a range of host barriers, including different defense systems, promises to provide further insights into how phage evolution occurs and the mechanisms responsible. In addition, such studies may also inform practical efforts to engineer phages and develop phage therapeutics.

## Acknowledgments

We thank A. Murray, K. Gozzi, I. Frumkin, C. Vassallo, and T. Zhang for comments on the manuscript, S. Jones and C. Eickmann for assistance in constructing plasmids, and all members of the Laub lab for helpful discussions. We thank the MIT BioMicro Center and its staff for their support in sequencing. This work was supported by an NSF predoctoral graduate fellowship to C.K.G. M.T.L. is an Investigator of the Howard Hughes Medical Institute.

## Author Contributions

S.S. performed phage evolution experiments, sequencing experiments, bioinformatic analyses, and phage growth and spotting experiments. C.K.G. performed biochemical experiments, cell growth assays, and phage growth and spotting experiments. S.S., C.K.G., and M.T.L. designed experiments, analyzed data, prepared figures, and wrote the manuscript.

## Resource Availability

### Lead Contact

Questions about or requests for methods, strains, and resources generated in this study can be directed to the Lead Contact, Michael T. Laub.

### Materials Availability

Plasmids and strains generated in this study are available upon request to the Lead Contact.

### Data and Code Availability

DNA sequencing data is available at SRA (BioProject ID: PRJNA824875). Scripts used for sequencing data processing and analysis are available at (https://github.com/sriramsrikant/2022_T4-toxIN-evo).

## Methods

### Strains and growth conditions

All strains, plasmids, and primers used in this study are listed in Tables S1, S2, and S3, respectively. For all phage experiments in liquid media and all phage spotting experiments, *Escherichia coli* MG1655 strains were grown in Luria broth (LB) medium. For pBAD33-ToxN induction experiments, cells were grown in M9 (10x stock made with 64 g/L Na_2_HPO_4_-7H_2_O, 15 g/L KH_2_PO_4_, 2.5 g/L NaCl, 5.0 g/L NH_4_Cl) medium supplemented with 0.1% casamino acids, 0.4% glycerol, 0.4% glucose, 2 mM MgSO_4_, and 0.1 mM CaCl_2_ (M9-glucose). For plasmid construction, *E. coli* DH5α cells were grown in Luria broth (LB) medium. Antibiotics were used at the following concentrations (liquid ; plates): carbenicillin (50 μg/mL ; 100 μg/mL), chloramphenicol (20 μg/mL ; 30 μg/mL).

### Plasmid construction

pKVS45-*tifA* and pKVS45-*dmd*: T4 *tifA* and *dmd* were individually PCR-amplified from purified T4 genomic DNA with primer pairs SS-5/SS-6 and SS-7/SS-8, respectively, which also add flanking BamHI and KpnI sites. These PCR products were then cloned into the BamHI and KpnI sites of the aTc-inducible pKVS45 vector (pKVS45-*tifA* and pKVS45-*dmd*). pKVS45-*tifA*-FLAG was constructed via round-the-horn PCR on pKVS45-*tifA* using the primer pair SS-9/10. pKVS45-*tifA* ΔATG was constructed via round-the-horn PCR on pKVS45-*tifA* using the primer pairs SS-31/32 followed by blunt-end ligation. pKVS45-*tifA* ablated ToxN-site was constructed via round-the-horn PCR on pKVS45-*tifA* using the primer pairs SS-33/34 followed by Gibson Assembly using the 2x HiFi DNA Assembly Master Mix (NEB E2621). TifA codons were recoded using Geneious and ordered as a gene-block fragment from IDT (with 46 of 85 codons recoded with 55 nucleotide changes) and cloned into pKVS45 by Gibson assembly at the same site as pKVS45-*tifA*. For pKVS45-*tifA*-homologs, *tifA* homologs from T2, T6 and RB69 were each PCR-amplified from purified phage genomic DNA with primer pairs SS-11/12, SS-13/14, and SS-15/16, respectively, which also add sequences that overlap the aTc-inducible pKVS45 vector. pKVS45 was digested with BamHI and KpnI, and then the PCR-amplified *tifA* variants were inserted into the digested plasmid using Gibson assembly.

pEXT20-*tifA*: The *tifA* homologs from T2, T4, T6 and RB69 were each PCR-amplified from purified phage genomic DNA with primer pairs SS-17/18, SS-19/20, SS-21/22 and SS-23/24, respectively, which also add sequences that overlap the IPTG-inducible pEXT20 vector. pEXT20 was digested with SacI and SalI, and then the PCR-amplified *tifA* variants were inserted into the digested plasmid using Gibson assembly.

pEXT20-*ipIII_T4_*: The *ipIII_T4_* gene was PCR amplified from T4 genomic DNA with primer pair SS-35/36 which add sequences that overlap the IPTG-inducible pEXT20 vector. pEXT20 was digested with SacI and SalI and *ipIII_T4_* was inserted into the digested plasmid using Gibson Assembly.

pBR322-*rIIA_T4_*: The *rIIA* gene including the upstream native promoter was PCR amplified from T4 genomic DNA with primer pair SS-37/38 which add sequences that overlap with the pBR322 vector. The pBR322 backbone was amplified with primer-pair SS-39/40, and the *rIIA* gene was inserted using Gibson assembly.

pKVS45-*rIIB_T4_*: The *rIIB* gene (only the coding sequence) was PCR amplified from T4 genomic DNA with primer pair SS-41/42 which add flanking BamHI and KpnI sites. This PCR product was then cloned into the BamHI and KpnI sites of the aTc-inducible pKVS45 vector.

pBR322-*Dsr1_ECOR17_*: The *Dsr1_ECOR17_* gene including the upstream promoter (300 bp) was PCR amplified from ECOR17 genomic DNA with primer pair SS-43/44 which add sequences that overlap with the pBR322 vector. The pBR322 backbone was amplified with primer-pair SS-39/40, and the *rIIA* gene was inserted using Gibson assembly.

pKVS45-*nrdC.5_T4_*: The *nrdC.5_T4_* gene (only the coding sequence) was PCR amplified from T4 genomic DNA with primer pair SS-45/46 which add flanking BamHI and KpnI sites. This PCR product was then cloned into the BamHI and KpnI sites of the aTc-inducible pKVS45 vector.

### Strain construction

*E. coli* MG1655 *attB_′″_::toxIN* and *E. coli* MG1655 *attB_′″_::toxIN(K55A)*: To construct these strains, *toxIN* or *toxIN(K55A)* was inserted into the *E. coli* MG1655 chromosome using the CRIM system (Haldimann and Wanner, 2001). The *toxIN* locus, along with its native promoter and transcription terminators, was PCR-amplified from pBR322-*toxIN* with the primer pair SS-25/26. The CRIM plasmid pAH150 was linearized using PCR with SS-27/28 to remove the *araBAD* promoter and *rrnB* T1 and *rrnB* T2 terminators. *toxIN* was then inserted into linearized pAH150 using Gibson assembly to yield pAH150-*toxIN*. pAH150-*toxIN(K55A)* was then constructed via round-the-horn PCR of pAH150-*toxIN* with SS-29/30. CRIM insertion of these plasmids into the *attB_′″_* locus of *E. coli* MG1655 was then performed as previously described (LeRoux et al., 2021) using the pINT-ts helper plasmid. Single insertions were confirmed by PCR and Sanger sequencing.

T4 mutants were generated using a CRISPR-Cas system for targeted mutagenesis described previously (Duong et al., 2020). Briefly, sequences for RNA guides to target Cas9-mediated cleavage were designed using the toolbox in Geneious Prime 2021.2.2 targeting *tifA* but nowhere else in the T4 genome. The guides were inserted into the pCas9 plasmid and tested for their ability to target the T4 genome by restricting T4 deficient in β-glucosyltransferase (*b-gt*) glucosylation. Escape plaques were isolated, and the tifA locus was sequenced by Sanger sequencing, to identify 2 unique clones that have a 98 bp deletion and a 5 bp insertion in *tifA* that disrupt its coding sequence (Figure S2B).

### Phage experiments in solid and liquid media

Phage spotting assays were conducted as described in (Guegler and Laub, 2021) and were conducted at least three times independently, with representative experiments shown in the figures. Briefly, phage stocks isolated from single plaques were propagated in *E. coli* MG1655 at 37 °C in LB. To titer phage, stocks were mixed with *E. coli* MG1655 and melted LB + 0.5% agar and spread on LB + 1.2% agar plates at 37 °C. For phage spotting assays, the bacterial strain of interest was mixed with LB + 0.5% agar and spread on an LB + 1.2% agar + antibiotic plate. For experiments in which expression of *tifA* was induced with anhydrous tetracycline (P*_tet_*-*tifA*), 100 ng/μL anhydrous tetracycline (aTc) was added to LB + 0.5% agar at a final concentration of 100 ng/mL prior to mixing with bacterial cells. Phage stocks were then serially diluted in 1x FM buffer (20 mM Tris-HCl pH 7.4, 100 mM NaCl, 10 mM MgSO_4_), and 2-3 μL of each dilution was spotted on the bacterial lawn. Plates were then grown at room temperature or 37 °C overnight and plaques quantified the following day. Efficiency of plaquing (EOP) was calculated by comparing the ability of the phage to form plaques on an experimental strain relative to the control strain.

Phage experiments in liquid culture were conducted at 37 °C. Single colonies were grown overnight in LB. Overnight cultures were back-diluted to OD_600_ = 0.01 and grown to OD_600_ = 0.3 in fresh LB such that cells re-entered exponential growth. Cultures were then back-diluted to OD_600_ = 0.01 in fresh LB, and phage was added to each culture. Growth following phage infection was measured in 24-well plates at 15 min intervals with orbital shaking at 37 °C on a plate reader (Biotek). Data points reported are the mean of two technical replicates each for three independent growth curve experiments.

### High-throughput evolution of T4 to infect resistant *E. coli*

Phage populations are evolved to infect novel hosts using a modification of the Appelmans protocol originally developed to generate phage therapy cocktails. This protocol was more recently used to modify the host range of phage (Burrowes et al., 2019; Mapes et al., 2016). Briefly, a population of phage is propagated on a sensitive host to maintain phage population size and generate genetic diversity, and also a resistant host to select for phage that can infect the novel host. In our experiment the resistant host was MG1655 with a *toxIN*-containing plasmid (ML3328), and the sensitive hosts are MG1655 with an empty plasmid (ML3330). Thus, T4 needs to successfully overcome the defense conferred by toxIN to successfully invade the resistant host. To visually track the progress of evolution, the evolution experiment was setup in 96-well plates with phage populations inoculated with host cells at a range of MOI (Supp. Figure 1). A phage blank well was also maintained to confirm there is no cross-contamination during the setup of each round of evolution.

T4 (10^6^ pfu/µL) lysate prepared in MG1655 was then serial diluted 10-fold in FM buffer to generate inoculum samples from 10^6^ - 1 pfu/µL. 10 µL of each the phage inoculums were added to the corresponding wells in a row. Each well contains 100 µL of supplemented LB (LB + 15 µg/mL L-tryptophan + 5 µg/mL thiamine) with 10^5^ bacterial cells. Bacterial culture was back-diluted from overnight growth of single colonies in LB (or M9) at 37 °C, using OD600 of 1 as an estimated cell density of 7 x 10^8^ cells/mL. The bacterial cells are ML3328 (*+toxIN*) and ML3330 (*-toxIN*) in two separate rows. The first wells of each row are inoculated with only 10 µL of FM buffer, as a no-phage control check against phage cross-contamination during the setup of evolution plate. The second to eighth wells of each row receive inoculum from the phage serial dilutions to setup wells with phage-to-bacteria rations (multiplicity of infection, MOI) of 10^2^ to 10^-4^. Each pair of rows correspond to an independent evolving population, with the top two rows in the plate being a control evolution with T4 infecting two rows of ML3330 to identify mutations that arise when T4 is passaged on just MG1655 with no *toxIN*. The remaining rows on the plate accommodate 5 independent evolving populations of T4. For the first round, the same clonal population of T4 is used to start all the evolving populations. After inoculating the cell culture with phage, the evolution plate is sealed with breathable film and incubated at 37 °C on a plate shaker at 1000 RPM overnight (14-18 hours). The plate is harvested by pooling all cleared wells and the first uncleared well (up to a total of 5 wells per row from the highest MOI) in every pair of rows to generate the population of phage at the end of the round. In a round of evolution, phage in the highest MOI well lyse all cells in a single infection generation, while at the lowest MOI (1 phage particle inoculated into 10^5^ bacteria) about 3 rounds of phage infection (with an estimated T4 burst of 100) are needed to lyse all the bacteria. Thus, a phage population after a round of evolution contains particles that have gone through 1-3 infection generations from a phage at the start of the round. The cell debris and unlysed cells in pooled samples were pelleted (4000RPM at 4 °C for 25 min) and the supernatant phage lysate was transferred to 96 deep-well blocks for long-term storage (with 40 µL chloroform to prevent bacterial growth). Pooled samples generated 6 populations - 5 replicate populations evolving on ML3328 and ML3330, and 1 control population evolving on ML3330 alone. Twenty-five rounds of evolution were performed saving a total of 150 populations (6 populations at each of 25 rounds of evolution). The evolution was performed in LB for the first 18 rounds and was then switched to M9 for 7 rounds to increase the selective pressure of *toxIN* on T4 replication. Evolving populations were stored as lysates at 4 °C, and also as glycerol stocks of infected hosts snap-frozen and left at −80 °C.

Evolved clones were isolated by plating to single plaques from the evolved populations on lawns of ML3328 for T4 that can overcome *toxIN*. These evolved clones were saved as phage lysates at 4 °C and as glycerol stocks of infected hosts for long-term storage.

### ToxN and TifA overexpression plating assays on solid media

Overexpression plating assays were conducted as described in (Guegler and Laub, 2021). Briefly, single colonies were grown to saturation overnight in M9-glucose. 1 mL of each overnight culture was then pelleted by centrifugation at 4000 *g* for 5 min, washed twice in 1x phosphate-buffered saline (PBS), and resuspended in 500 μL 1x PBS. Cultures were then serially-diluted 10-fold in 1x PBS and spotted on M9L plates (M9-glycerol supplemented with 5% LB (v/v)) further supplemented with 0.4% glucose (toxin-repressing), 0.2% arabinose (toxin inducing), or 0.2% arabinose and 100 μM IPTG (toxin and antitoxin inducing). Plates were then incubated at 37 °C for 48 h before imaging. Images shown are representative of at least three independent biological replicates.

### ToxN and TifA overexpression growth curves in liquid culture

Single colonies of *E. coli* MG1655 harboring pBAD33-*toxN* and pEXT20-*tifA_T4_* were grown to saturation overnight in M9-glucose. These cultures were then back-diluted 100-fold in fresh M9-glycerol supplemented with 0.8% glucose and grown to OD_600_ ∼0.3 at 37 °C in a shaking water bath; additional glucose was added to these cultures to suppress leaky toxicity of ToxN. Cells were pelleted by centrifugation at 4 °C and 4000 *g* for 5 minutes. Pellets were washed twice in ice-cold M9-glycerol and then resuspended in fresh M9-glycerol. Cells were then back-diluted in fresh M9-glycerol to OD_600_ = 0.2; for cultures in which TifA expression was induced, 100 μM IPTG was also added at this point. Cultures were then grown for an additional 45 min at 37 °C in a shaking water bath before being back-diluted to OD_600_ = 0.1 in fresh M9-glycerol. To these back-diluted cultures, 100 μM IPTG and 0.2% arabinose added to induce TifA and ToxN, respectively, when appropriate. Growth was then measured in 24-well plates at 15 min intervals with orbital shaking at 37 °C on a plate reader (Biotek). Data reported are the mean of three technical replicates each of four independent growth curve experiments.

### ToxN-TifA interaction assays with co-immunoprecipitation (co-IP)

50 mL cultures of *E. coli* MG1655 containing pBR322-*toxIN*(+/-His_6_) and pKVS45-*tifA*(+/- FLAG) were grown to OD_600_ = 0.4 at 37 °C in LB supplemented with 15 μg/mL L-tryptophan and 1 μg/mL thiamine in a water bath shaker and infected with T4 at MOI = 5. To induce expression of TifA, aTc was added to each culture at a final concentration of 100 ng/mL 30 min prior to phage infection. 15 min post-T4 infection, 40 mL of cells from each culture were pelleted by centrifugation at 4000 *g* for 5 min at 4 °C. Pellets were then flash-frozen and stored at −80 °C.

For anti-FLAG co-IP experiments, each cell pellet was resuspended in 500 μL of lysis buffer (25 mM Tris-HCl pH 8, 150 mM NaCl, 1 mM EDTA pH 8, 1% Triton X-100, cOmplete protease inhibitor cocktail (Roche), 1 μL/mL ReadyLyse (Novagen), 1 μL/mL DNaseI (TakaraBio), 5% glycerol). Cells were lysed by two freeze-thaw cycles, followed by incubation at 30 °C for 1 hr. Lysates were clarified by centrifugation at 14,000 *g* for 10 min at 4 °C and then incubated with Anti-FLAG-M2 magnetic beads (Sigma) on an end-to-end rotor for 1 hr at 4 °C. Beads were then washed three times with 400 μL wash buffer (25 mM Tris-HCl pH 8, 150 mM NaCl, 1 mM EDTA pH 8, cOmplete protease inhibitor cocktail (Roche), 5% glycerol). To elute protein, beads were incubated with 40 μL of 1x Laemmli buffer, vortexed briefly, and boiled at 95 °C for 5 min.

Samples from co-IP experiments were analyzed by Western blot as described previously (Guegler and Laub 2021). Briefly, samples were resolved by 4-20% SDS-PAGE and transferred to a PVDF membrane. Membranes were then incubated with rabbit anti-His_6_ and anti-FLAG antibodies (from Abcam and Cell Signaling Technologies, respectively) at a final concentration of 1:5000 to avoid cross-reactivity with the anti-FLAG-M2 beads. Blots were developed using the SuperSignal West Femto Maximum Sensitivity Substrate (Thermofisher) and imaged with a FluorChem R Imager (ProteinSimple). Blots shown are representative of three independent replicates.

### Phage genome isolation, Illumina library preparation and analysis

Phage genomes were prepared by treating high titer lysates (> 10^6^ pfu/µL) with DNAse I (0.001 U/µL) and RNAse A (0.05 mg/mL) at 37 °C for 30 min to hydrolyze any host nucleic acid that remains in the lysate. EDTA is added to a final of 10 mM to inactivate nucleases. The lysate is then incubated with Proteinase K at 50 °C for 30 min to disrupt phage capsids and release genomes. The phage genomes are recovered by Ethanol precipitation. Briefly, NaAc pH 5.2 was added to 300 mM followed by 100% ethanol to a final volume fraction of 70%. The samples were left at - 80 °C for 2 h to precipitate genomic material. The genomic material was pelleted at 21000 xg for 15 min, and the supernatant is discarded. The pellet was washed in series with 100 µL isopropanol, and 200 µL 70% (v/v) ethanol by pelleting and discarding supernatants for each wash. The pellet was air-dried at room temperature for 30 min and then resuspended in 20-100 µL TE (10mM Tris-HCl, 0.1mM EDTA, pH8) overnight at room temperature. Genomic material was resuspended by flicking the eppendorf and concentration was measured using a Nanodrop spectrophotometer.

Illumina sequencing libraries were prepared starting from 100-200 ng of genomic DNA (gDNA) as described before. Briefly, the gDNA was sheared by sonication in a Bioruptor. Fragmented gDNA was purified using AmpureXP beads, followed by sequential enzymatic reactions for end repair, 3’ adenylation, and adapter ligation. Barcodes were added to both 5’ and 3’ ends by PCR amplifying with primers that anneal to the Illumina PE adapters. The libraries were cleaned and by Ampure XP clean up using a double cut to elute fragment sizes matching the read-lengths of the sequencing run. Libraries were sequenced on either an Illumina MiSeq or NextSeq at the MIT BioMicroCenter, after qPCR and fragment analyzer quality control for concentration and size distribution.

Illumina reads were analyzed using custom python scripts. Briefly, the reads were cleaned of adapter sequences using cutadapt before they are aligned to a reference genome using bwa and samtools. Per position coverage maps were generated using samtools depth and variant single-nucleotide polymorphisms were called using bcftools. Coverage maps in figures were plotted by using a moving window average of per nucleotide coverage, with the window size defined as the read length of sequencing run. The ancestral T4 was sequenced to generate a reference genome against which all evolved clones were compared. Illumina reads were also assembled to reference genomes using the assembly pipeline in Geneious Prime 2021.2.2 for cross-checks and visual inspection.

### Homology search and alignment of TifA

Homologs of TifA were first identified by jackhmmer using a Hidden Markov Model built on a small subset of TifA homologs among reference proteomes on UniProt. The overwhelming majority of homologs with full query coverage were in viruses that are members of the phage clade Caudovirales. These 57 sequences were aligned using MUSCLE (with default settings), and a maximum likelihood phylogenetic tree was constructed with FastTree. The Tree separated TifA homologs from different phage genomes into clusters by similarity (color coded, Figure S3).

---

**Table S1.** Strains.

| Bacterial Strains |  |  |
| --- | --- | --- |
| Name | Genotype | Source |
| ML6 | MG1655 |  |
| DH5α | Cloning strain | Invitrogen |
| ML3326 | MG1655 pBAD33-*toxN* pEXT20-*toxI* | Guegler and Laub, 2021 |
| ML3328 | MG1655 pBR322-*toxIN* | Guegler and Laub, 2021 |
| ML3330 | MG1655 pBR322 | Guegler and Laub, 2021 |
| ML3772 | MG1655 pBR322-*toxIN* pKVS45 | This study |
| ML3773 | MG1655 pBR322-*toxIN* pKVS45-*dmd*<sub>T4</sub> | This study |
| ML3774 | MG1655 pBR322-*toxIN* pKVS45-*tifA*<sub>T4</sub> | This study |
| ML3775 | MG1655 pBR322-*toxIN* pKVS45-*tifA*<sub>T2</sub> | This study |
| ML3776 | MG1655 pBR322-*toxIN* pKVS45-*tifA*<sub>T6</sub> | This study |
| ML3777 | MG1655 pBR322-*toxIN* pKVS45-*tifA*<sub>RB69</sub> | This study |
| ML3778 | MG1655 pBAD33-*toxN* pEXT20-*tifA*<sub>T2</sub> | This study |
| ML3779 | MG1655 pBAD33-*toxN* pEXT20-*tifA*<sub>T4</sub> | This study |
| ML3780 | MG1655 pBAD33-*toxN* pEXT20-*tifA*<sub>T6</sub> | This study |
| ML3781 | MG1655 pBAD33-*toxN* pEXT20-*tifA*<sub>RB69</sub> | This study |
| ML3782 | MG1655 pBAD33-*toxN* pEXT20 | This study |
| ML3783 | MG1655 pBR322-*toxIN* pKVS45-*tifA*<sub>T4</sub> ΔATG | This study |
| ML3784 | MG1655 pBR322-*toxIN* pKVS45-*tifA*<sub>T4</sub> recoded | This study |
| ML3785 | MG1655 pBR322-*toxIN* pKVS45-*tifA*<sub>T4</sub> ablated *toxN*-site | This study |
| ML3786 | MG1655 pBR322-*toxI*-*toxN*-His<sub>6</sub> pKVS45-*tifA*<sub>T4</sub>-FLAG | This study |
| ML3787 | MG1655 pBR322- *toxI*-*toxN*-His<sub>6</sub> pKVS45-*tifA*<sub>T4</sub> | This study |
| ML3788 | MG1655 pBR322-*toxI*-*toxN* pKVS45-*tifA*<sub>T4</sub>-FLAG | This study |
| ML3789 | MG1655 *attB*<sub>λ</sub>:*toxIN* | This study |
| ML3790 | MG1655 *attB*<sub>λ</sub>:*toxI*-*toxN*(K55A) | This study |
| ML3791 | MG1655 λ-lysogen | This study |
| ML3792 | MG1655 λ-lysogen pBR322-*rIIA*<sub>T4</sub> pKVS45-*rIIB*<sub>T4</sub> | This study |
| ML3793 | *E. coli* str C | Félix d'Hérelle<br>Reference Center for<br>Bacterial Viruses,<br>Université Laval |
| ML3794 | ECOR17 | Thomas S. Whittam<br>STEC Center at<br>Michigan State<br>University |
| ML3795 | ECOR17 pEXT20-*ipIII*<sub>T4</sub> | This study |
| ML3796 | ECOR17 pEXT20-*ipIII*<sub>T4</sub>ΔCTS | This study |
| ML3797 | ECOR17 ΔRM-*typeI* | This study |
| ML3798 | ECOR17 ΔRM-*typeIII* | This study |
| ML3799 | ECOR17 Δabi2 | This study |
| ML3800 | ECOR17 $\Delta dsrI$ | This study |
| ML3801 | ECOR17 $\Delta hhe$ | This study |
| ML3802 | ECOR17 $\Delta cas3$ | This study |
| ML3803 | ECOR71 | Thomas S. Whittam<br>STEC Center at<br>Michigan State<br>University |
| ML3804 | ECOR13 | Thomas S. Whittam<br>STEC Center at<br>Michigan State<br>University |
| ML3805 | ECOR16 | Thomas S. Whittam<br>STEC Center at<br>Michigan State<br>University |
| ML3806 | MG1655 pBR322-*DsrI*<sub>ECOR17</sub> pKVS45 | This study |
| ML3807 | MG1655 pBR322-*DsrI*<sub>ECOR17</sub> pKVS45-*nrdC.5*<sub>T4</sub> | This study |
| ML3808 | MG1655 pCas9-*tifA*<sub>T4</sub>-cr4 | This study |
| ML3343 | DH5α pBAD33-*toxN* | Guegler and Laub, 2021 |
| ML3345 | DH5α pEXT20-*toxI* | Guegler and Laub, 2021 |
| ML1978 | DH5α pEXT20 | E. coli Genetic Stock<br>Center, #12325 |
| ML3346 | DH5α pBR322-*toxIN* | Guegler and Laub, 2021 |
| ML3348 | DH5α pBR322 | Guegler and Laub, 2021 |
| ML3349 | DH5α pBR322-*toxI*-*toxN*-His<sub>6</sub> | Guegler and Laub, 2021 |
| ML3809 | DH5α pKVS45-*dmd*<sub>T4</sub> | This study |
| ML3810 | DH5α pKVS45-*tifA*<sub>T4</sub> | This study |
| ML3811 | DH5α pKVS45-*tifA*<sub>T2</sub> | This study |
| ML3812 | DH5α pKVS45-*tifA*<sub>T6</sub> | This study |
| ML3813 | DH5α pKVS45-*tifA*<sub>RB69</sub> | This study |
| ML3814 | TOP10 pEXT20-*tifA*<sub>T2</sub> | This study |
| ML3815 | TOP10 pEXT20-*tifA*<sub>T4</sub> | This study |
| ML3816 | TOP10 pEXT20-*tifA*<sub>T6</sub> | This study |
| ML3817 | TOP10 pEXT20-*tifA*<sub>RB69</sub> | This study |
| ML3818 | DH5α pKVS45-*tifA*<sub>T4</sub> ΔATG | This study |
| ML3819 | DH5α pKVS45-*tifA*<sub>T4</sub> recoded | This study |
| ML3820 | DH5α pKVS45-*tifA*<sub>T4</sub> ablated *toxN*-site | This study |
| ML3821 | TOP10 pKVS45-*tifA*<sub>T4</sub>-FLAG | This study |
| ML3822 | DH5α pBR322-*rIIA*<sub>T4</sub> | This study |
| ML3823 | DH5α pKVS45-*rIIB*<sub>T4</sub> | This study |
| ML3824 | DH5α pEXT20-*ipIII*<sub>T4</sub> | This study |
| ML3825 | DH5α pEXT20-*ipIII*<sub>T4</sub>ΔCTS | This study |
| ML3826 | DH5α pBR322-*DsrI*<sub>ECOR17</sub> | This study |
| ML3827 | DH5α pKVS45-*nrdC.5*<sub>T4</sub> | This study |
| ML3828 | DH5α pCas9-*tifA*<sub>T4</sub>-cr4 | This study |
| **Phage Strains** |  |  |
| **Name** | **Genotype** | **Source** |
| phML31 | T4 ancestor | Guegler and Laub, 2021 |
| phML32 | T4 control evo round 25 clone 1 | This study |
| phML33 | T4 evo 1 round 25 clone 1 | This study |
| phML34 | T4 evo 2 round 25 clone 1 | This study |
| phML35 | T4 evo 3 round 25 clone 1 | This study |
| phML36 | T4 evo 4 round 25 clone 1 | This study |
| phML37 | T4 evo 5 round 25 clone 1 | This study |
| phML38 | T2 | ATCC Cat #: 11303-B5 |
| phML39 | T6 | ATCC Cat #: 11303-B5 |
| phML40 | RB69 | Laval Collection, HER # 158 |
| phML41 | T4 *tifA-1* | This study |
| phML42 | T4 *tifA-2* | This study |

**Table S2.**
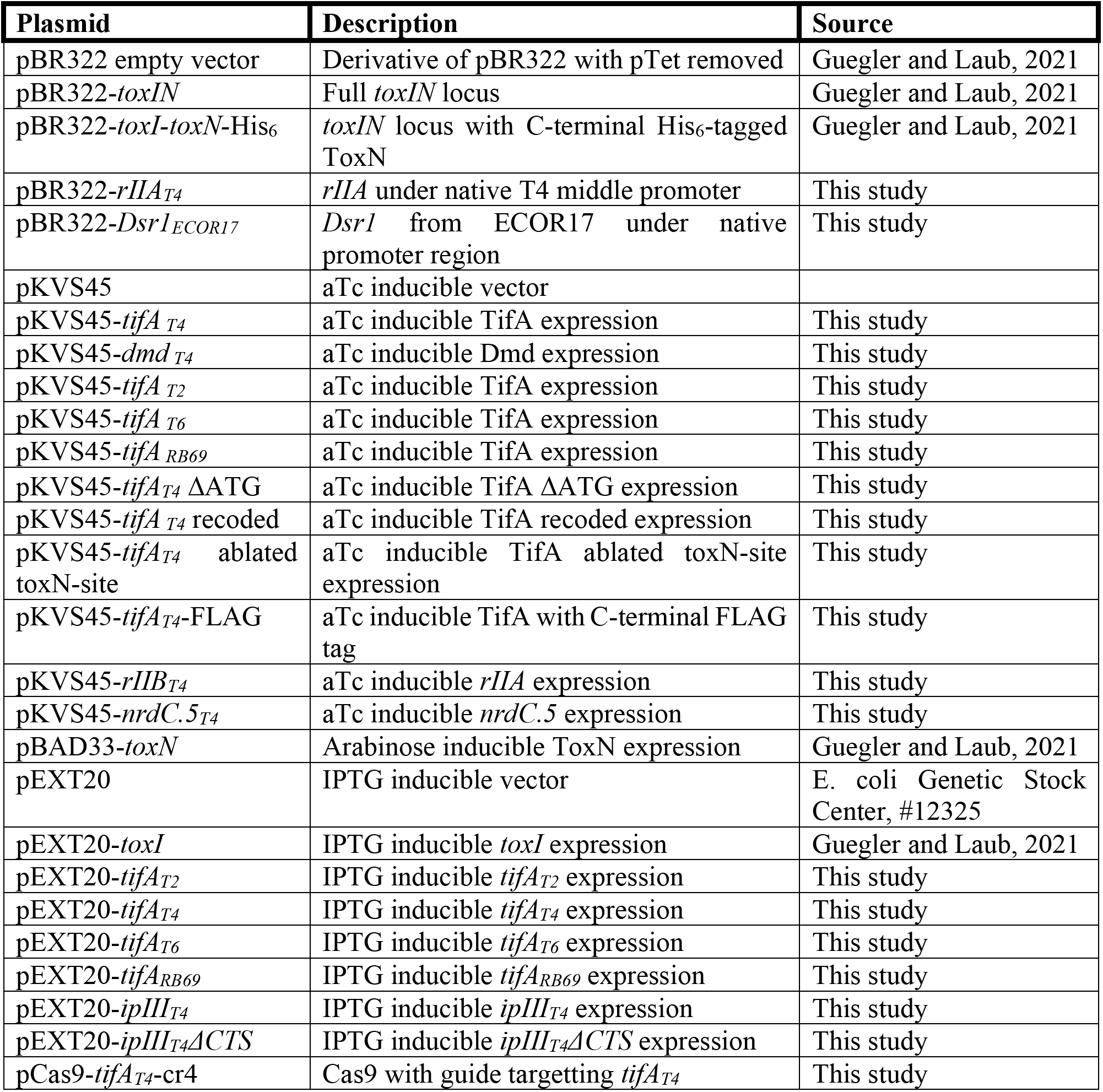
Plasmids.

**Table S3.**
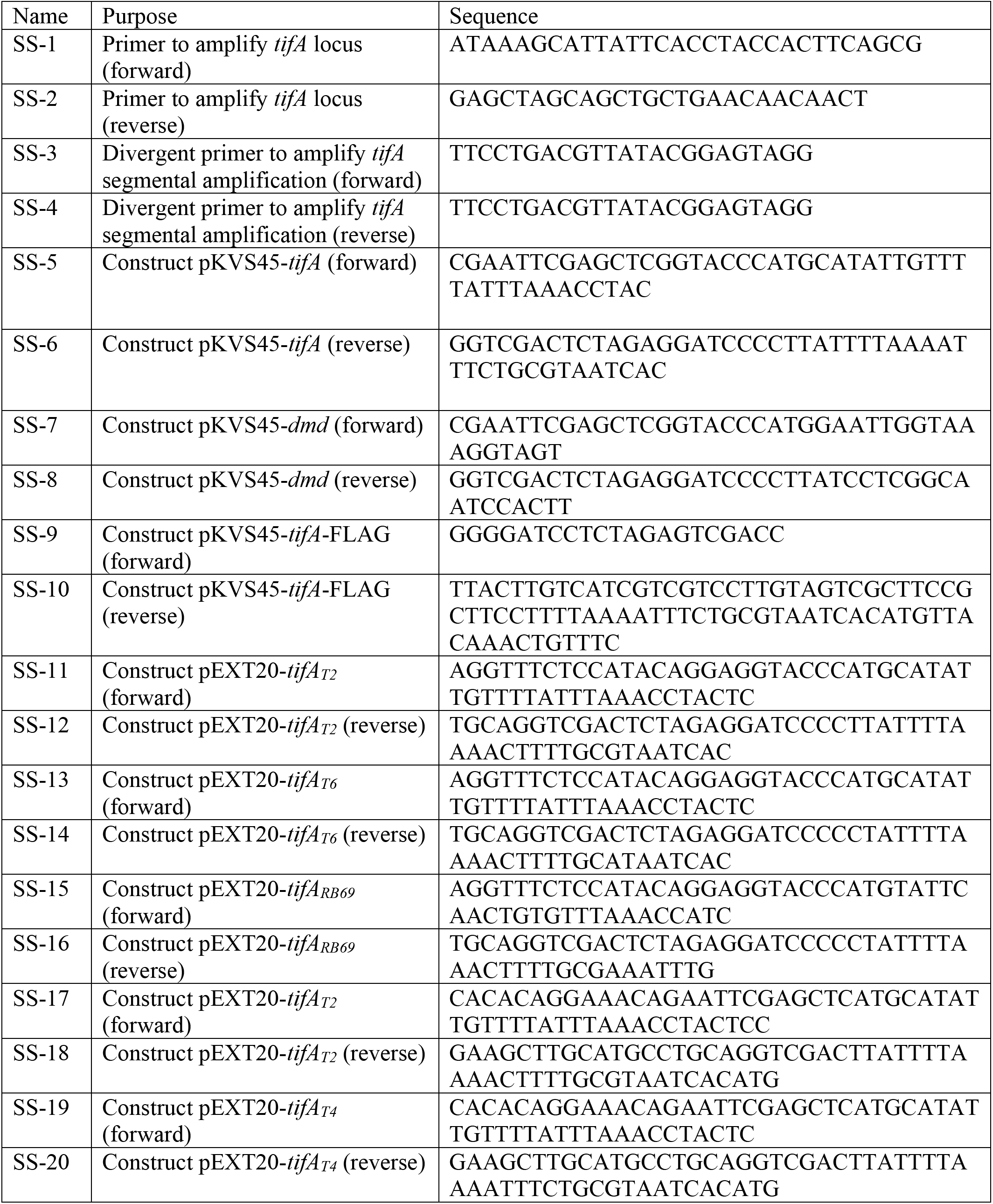
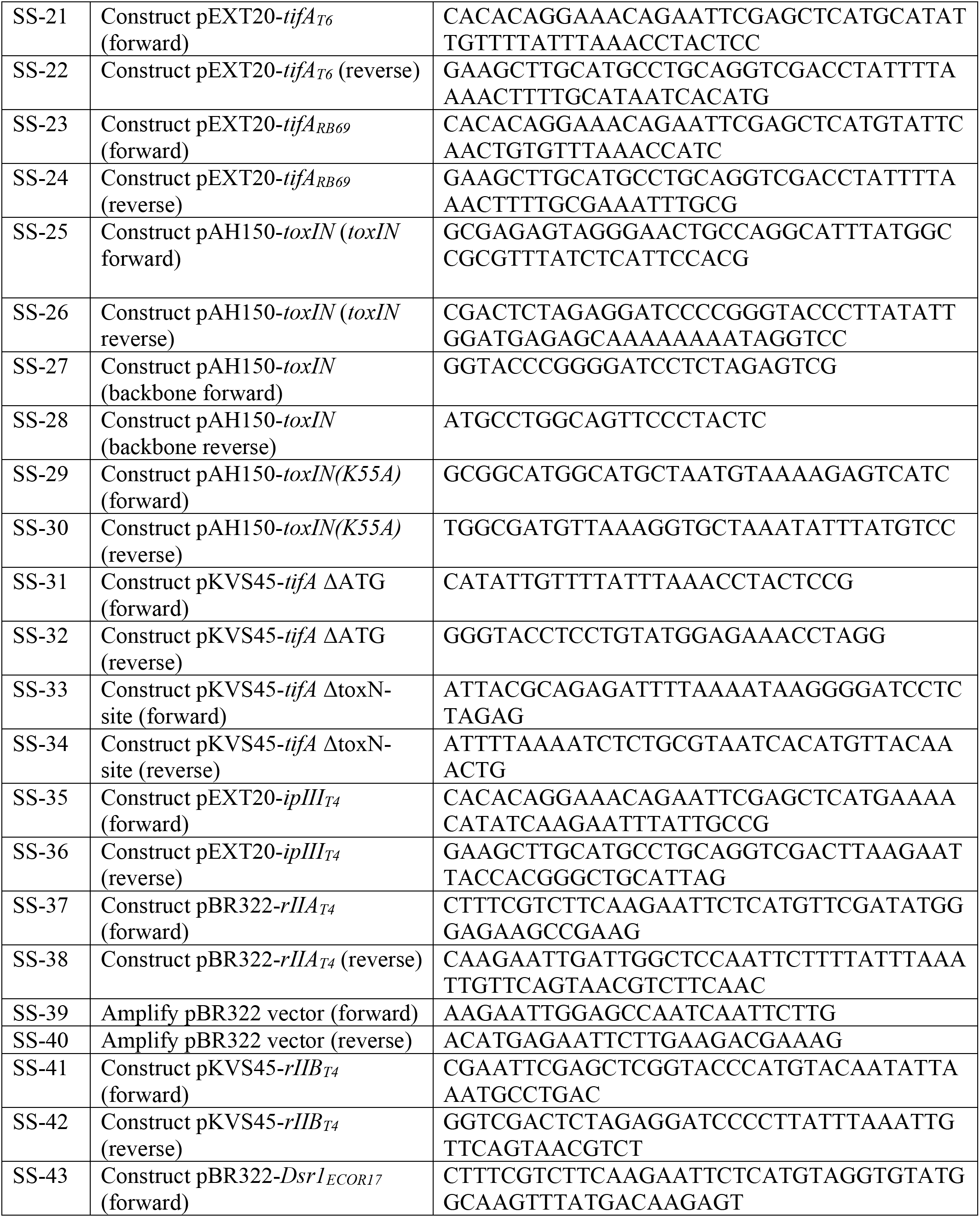
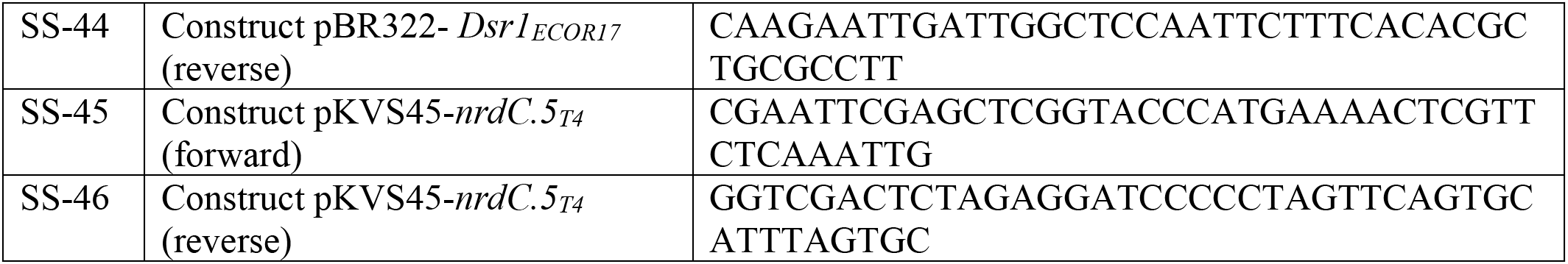
Primers.

**Figure S1.**
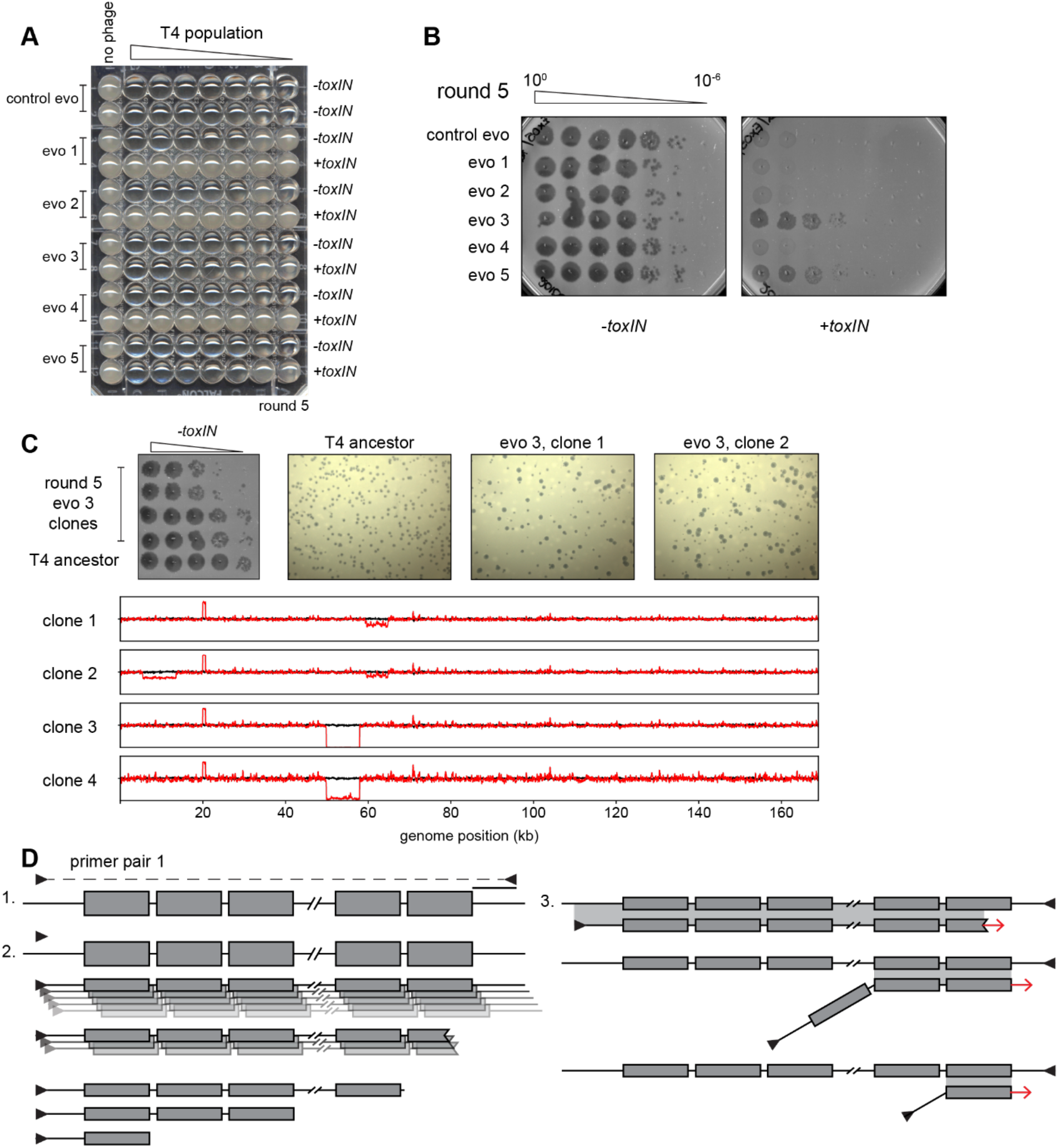
Experimental evolution of T4 to overcome *toxIN*. (**A**) Experimental evolutions were performed in 96-well plates. Pairs of rows were used for each evolving population and contain *- toxIN* or *+toxIN* cells in consecutive rows except for the control evolution which has *-toxIN* cells in both rows. Plates were imaged after overnight growth with clearing in wells indicating successful phage replication. The plate shown corresponds to round 5. (**B**) Serial dilutions of evolving populations spotted on *-toxIN* (*left*) and *+toxIN* (*right*) cells after 5 rounds of evolution shown as an example of how population size and ability to infect *+toxIN* cells were monitored. (**C**) Serial dilutions (*top left*) and plaque assays (*top right*) of plaque clone isolates 1 and 2 from evo 3 after round 5 shows dramatic plaque size variability compared to the T4 ancestor. (*Below*) Fold coverage of evolved clones (red lines) compared to T4 ancestor (black line) highlighting genomic differences across clones and deletions that arise in sub-populations during amplification from a single plaque. (**D**) Schematic representing generation of laddered products from PCR amplification of template with tandem repeats (step 1). Polymerase extension from an annealed primer generates a complementary strand to the template of length depending on the processivity of the polymerase. With purified polymerases and sufficient extension time, most products cover a large part of the repeated region, but some events generate smaller fragments containing fewer repeats (step 2). These smaller fragments can serve as primers in subsequent rounds of amplification by efficiently annealing to the template, but at mismatched repeats (step 3). This generates all sizes of products differing in the number of repeats with flanking primer annealing sites.

**Figure S2.**
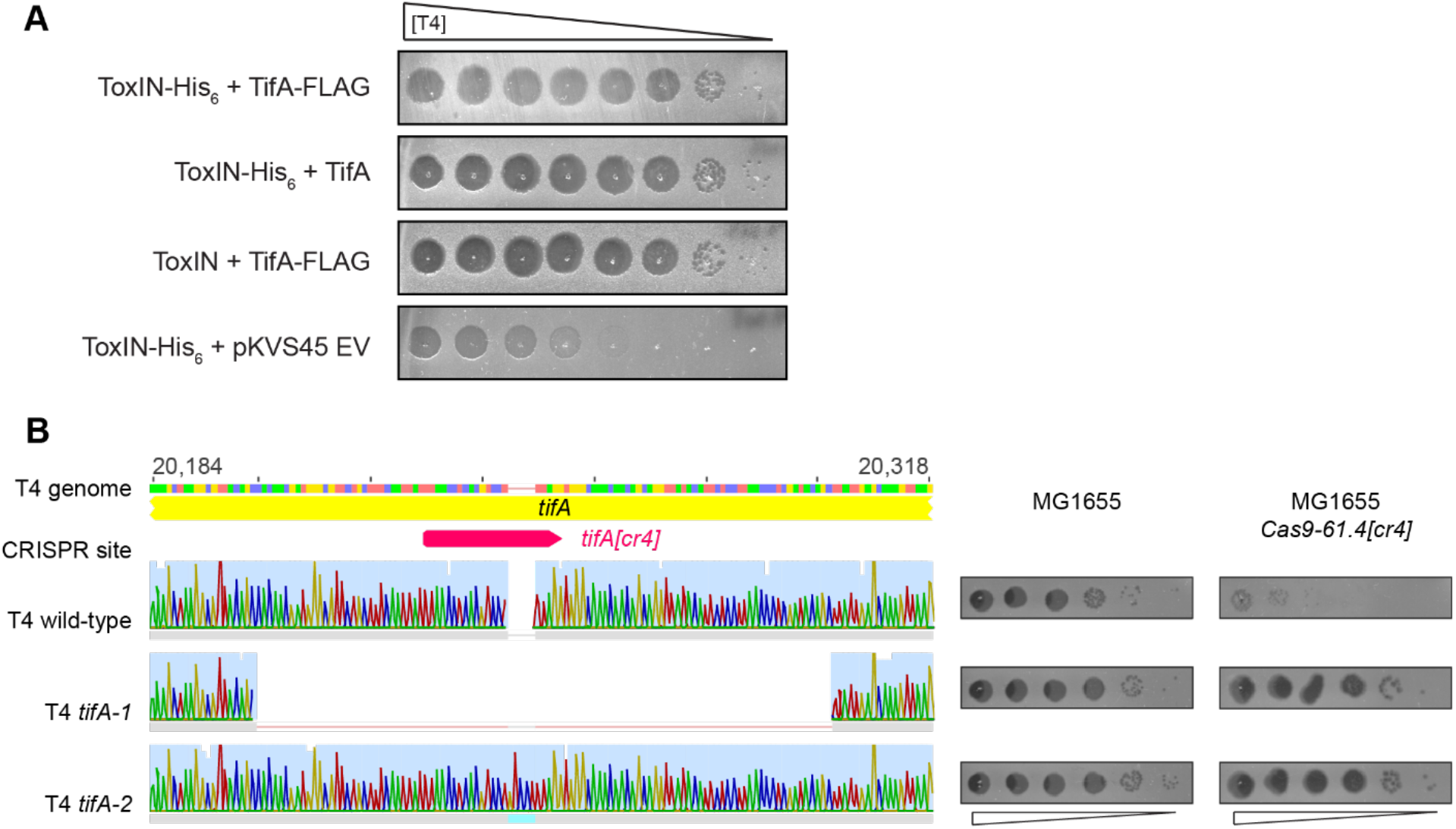
TifA-FLAG is functional in the context of T4 infection. (**A**) Serial dilutions of T4 ancestor on lawns of *E. coli* MG1655 containing *toxIN* or *toxI-toxN*-His_6_ and co-expressing *tifA* or *tifA*-FLAG, or carrying an empty vector, as indicated. (**B**) T4 mutant clones with mutations in *tifA*. (*Left*) Sanger sequencing chromatograms of *tifA* in wild-type T4, *tifA-1* (98 bp deletion), and *tifA-2* (5 bp insertion). (*Right*) Serial dilutions of T4 and *tifA* mutants on MG1655 and MG1655 expressing Cas9 with the *tifA[cr4]* guide-RNA shown above the chromatograms.

**Figure S3.**
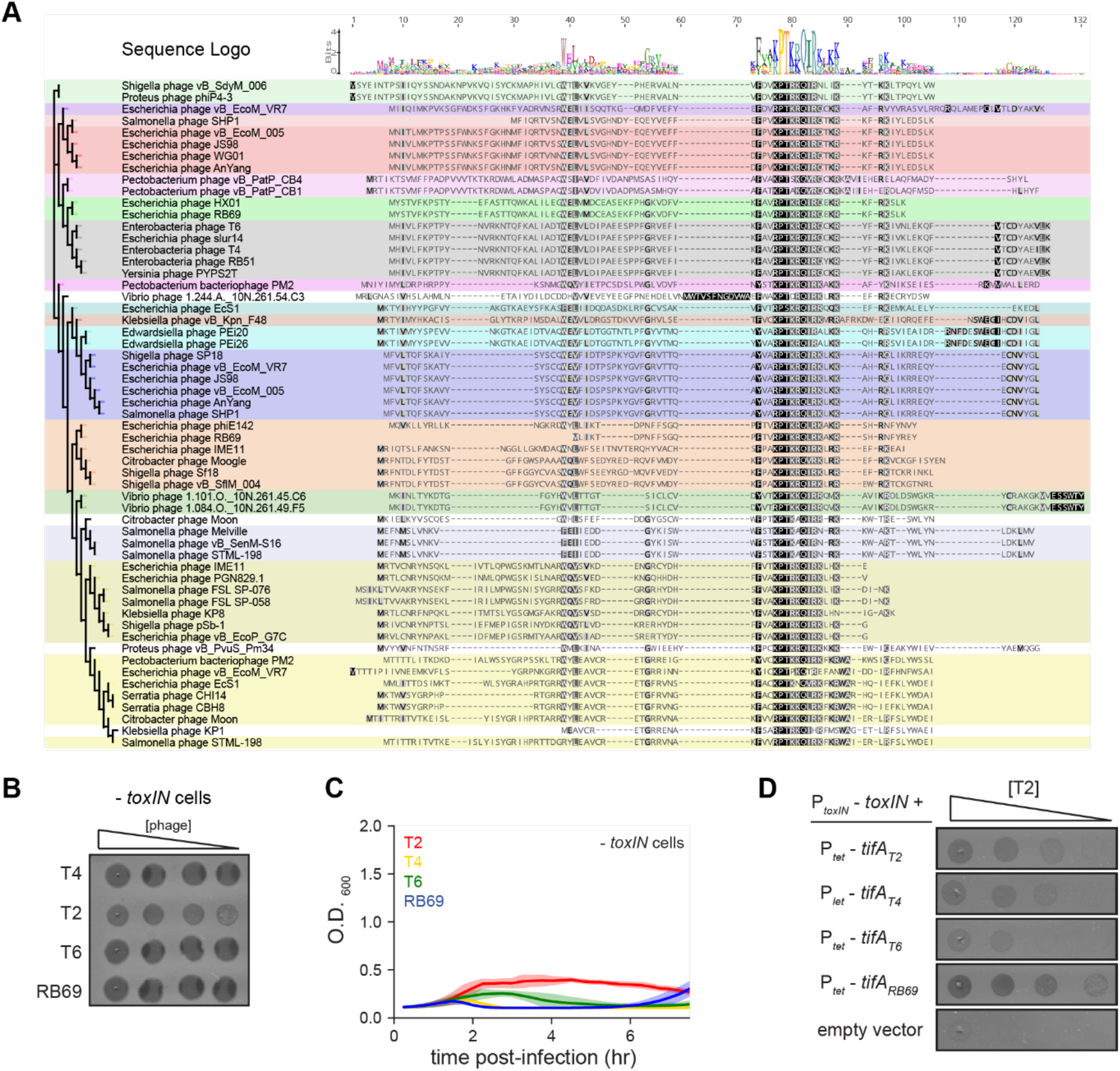
Homologs of *tifA* are found in other phage genomes. (**A**) Multiple sequence alignment of 57 TifA proteins found across sequenced phage genomes. TifA homologs have a conserved central region (highlighted by darker shades of grey) with varied N- and C-terminal regions. Groups of similar sequences, as judged by the tree (left), are shaded with the same color. Sequences were identified by jackhmmer, aligned by MUSCLE, and a pseudo-maximum likelihood tree built with FastTree. (**B**) Serial dilution plaquing assays of T4, T2, T6, and RB69 from Figure 3B on *-toxIN* cells. (**C**) Growth curves following infection of *-toxIN* cells with T4, T2, T6, or RB69, each at a MOI of 10^-3^. (D) Serial dilutions of T2 spotted on *+toxIN* cells expressing *tifA* homologs from T2, T4, T6, RB69, or harboring an empty vector.

**Figure S4.**
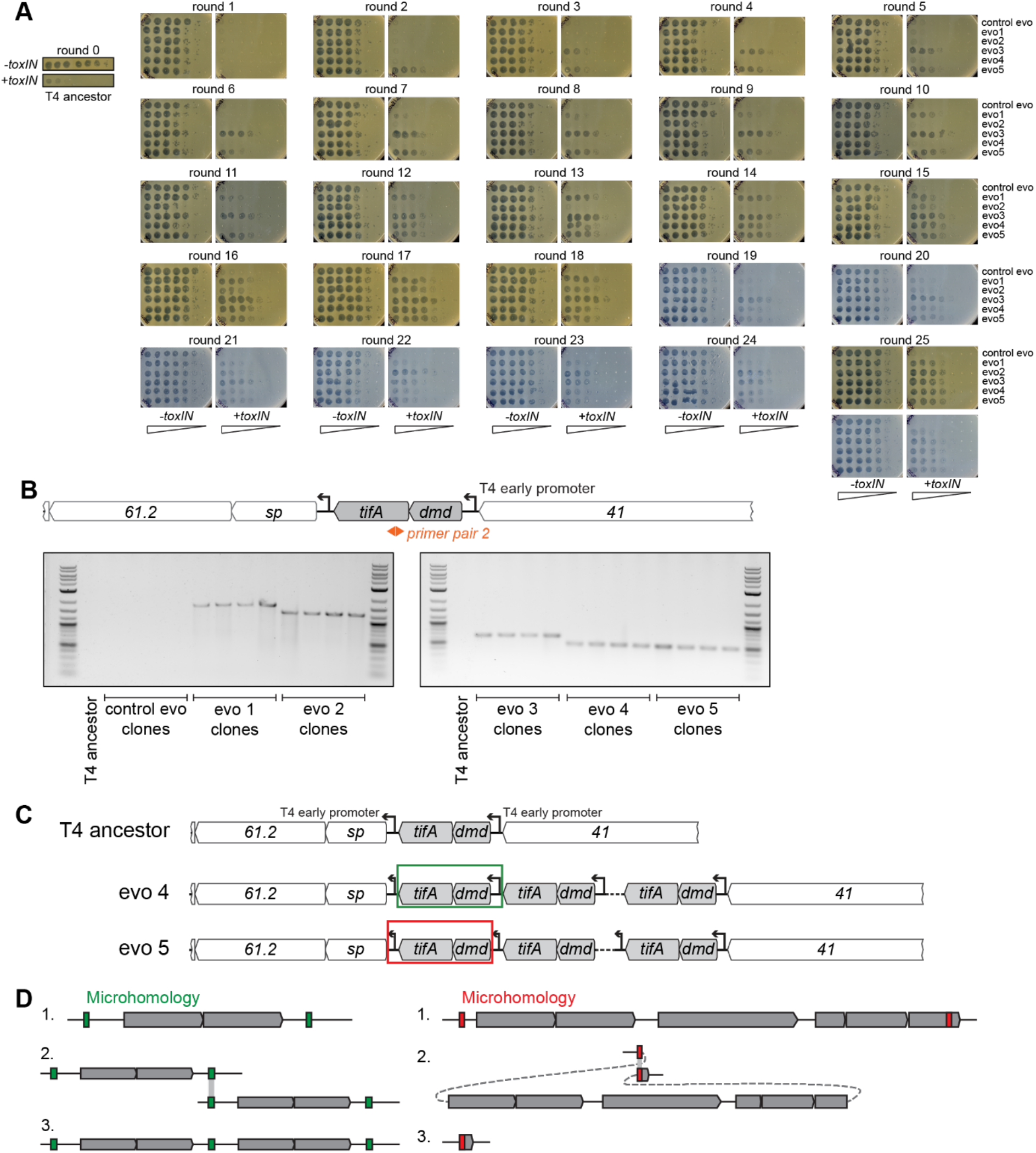
T4 replicate evolutions overcome *toxIN* using similar segmental amplifications of *tifA*. (**A**) Population size of replicate evolving T4 populations are determined by the titer estimated on *-toxIN* lawns and remain roughly constant (10^5^-10^6^ pfu/µL) over 25 rounds. Serial-dilutions of evolving populations (control evo, evo 1 to evo 5, top-to-bottom in each image) spotted on -*toxIN* and +*toxIN* lawns. These data were quantified to generate the heatmap in Figure 4B. (**B**) PCR products with divergent primers (primer pair 2, used similar to Figure 1) confirm presence of contiguous repeats in clones isolated from each evolving population (but not in clones from control evo or the T4 ancestor), with different sizes confirming unique repeating units in each evolving population’s amplification. (**C**) Schematic of repeats in evolved clones that link either the T4 early promoter upstream of *dmd-tifA* (evo 4, middle) or the T4 early promoter downstream of *dmd-tifA* (evo 5, bottom) to ensure expression of operon in all repeats. (**D**) Schematics of recombination events that can lead to segmental amplifications (*left*) or genome deletions (*right*). T4 is known to rely on recombination during genome replication at late times and is also used in the concatenation of genomes to enable processive genome packaging. Recombination between short identical sequence repeats, or regions of microhomology (green or red), across or within genome molecules can generate a segmental copy or loss of the intervening region, respectively.

**Figure S5.**
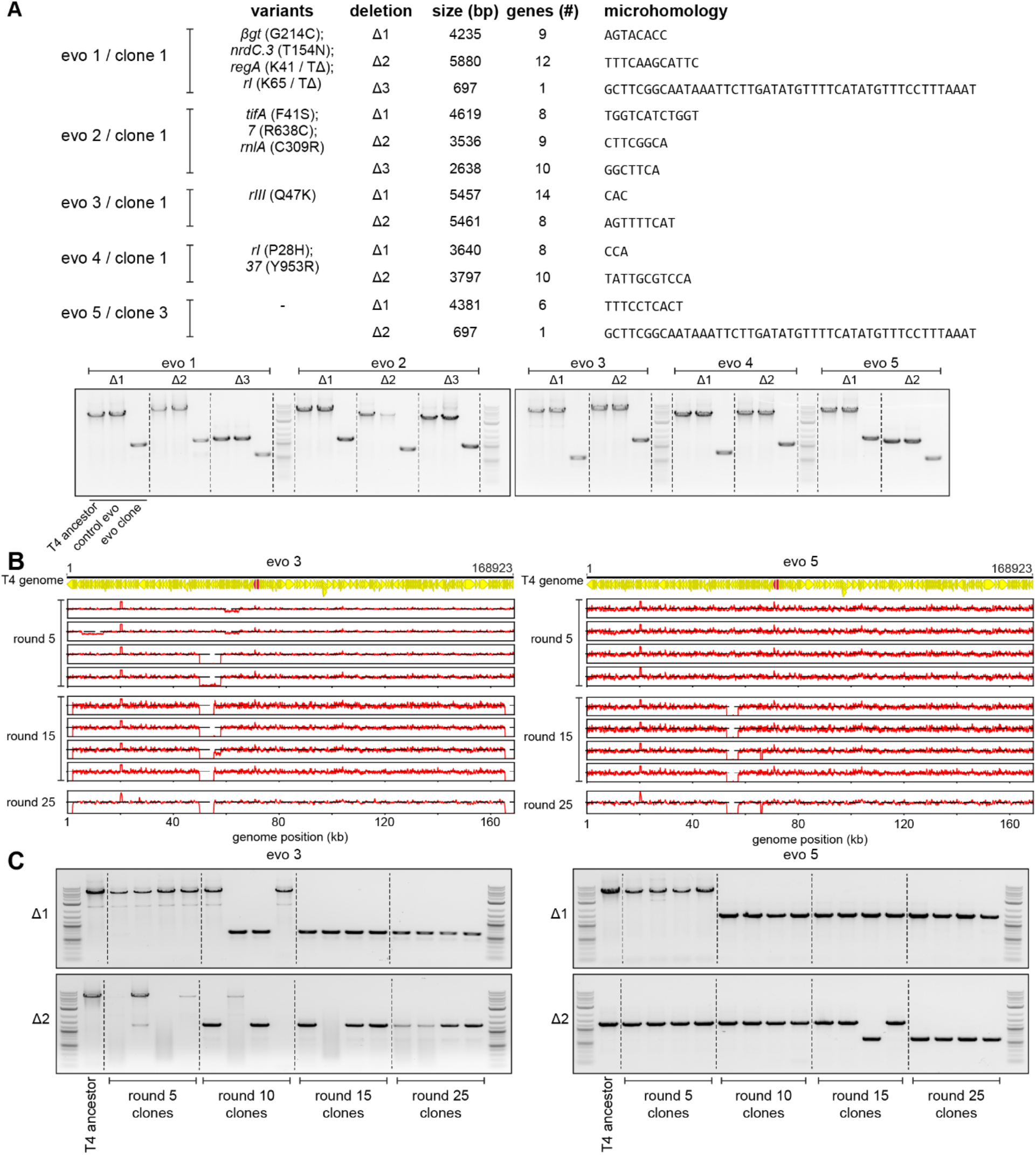
Genome deletions fix over the course of the evolution to compensate for amplifications at the *dmd-tifA* locus. (**A**) (Top) Summary of mutations and deletions identified by genome sequencing of isolated clones from each population after 25 rounds of evolution. Boundaries of the deletion, which are short regions of microhomology, are listed along with the deleted genes. (Bottom) PCR with primers flanking each of the deleted regions, to confirm smaller products generated from the corresponding evolved clone compared to T4 ancestor (or clone from control evo). (**B**) Fold coverage calculated from genome sequencing of clones isolated from populations 3 and 5 after various rounds of evolution (5, 15, 25) to follow amplifications and deletions that arise in evo 3 (*left*) and evo 5 (*right*). (**C**) PCR with primers flanking the deleted regions of evolving populations 3 (*left*) and 5 (*right*) performed on evolved clones isolated from rounds of evolution (5, 10, 15, 25) to trace genome configurations over evolutionary time.

**Figure S6.**
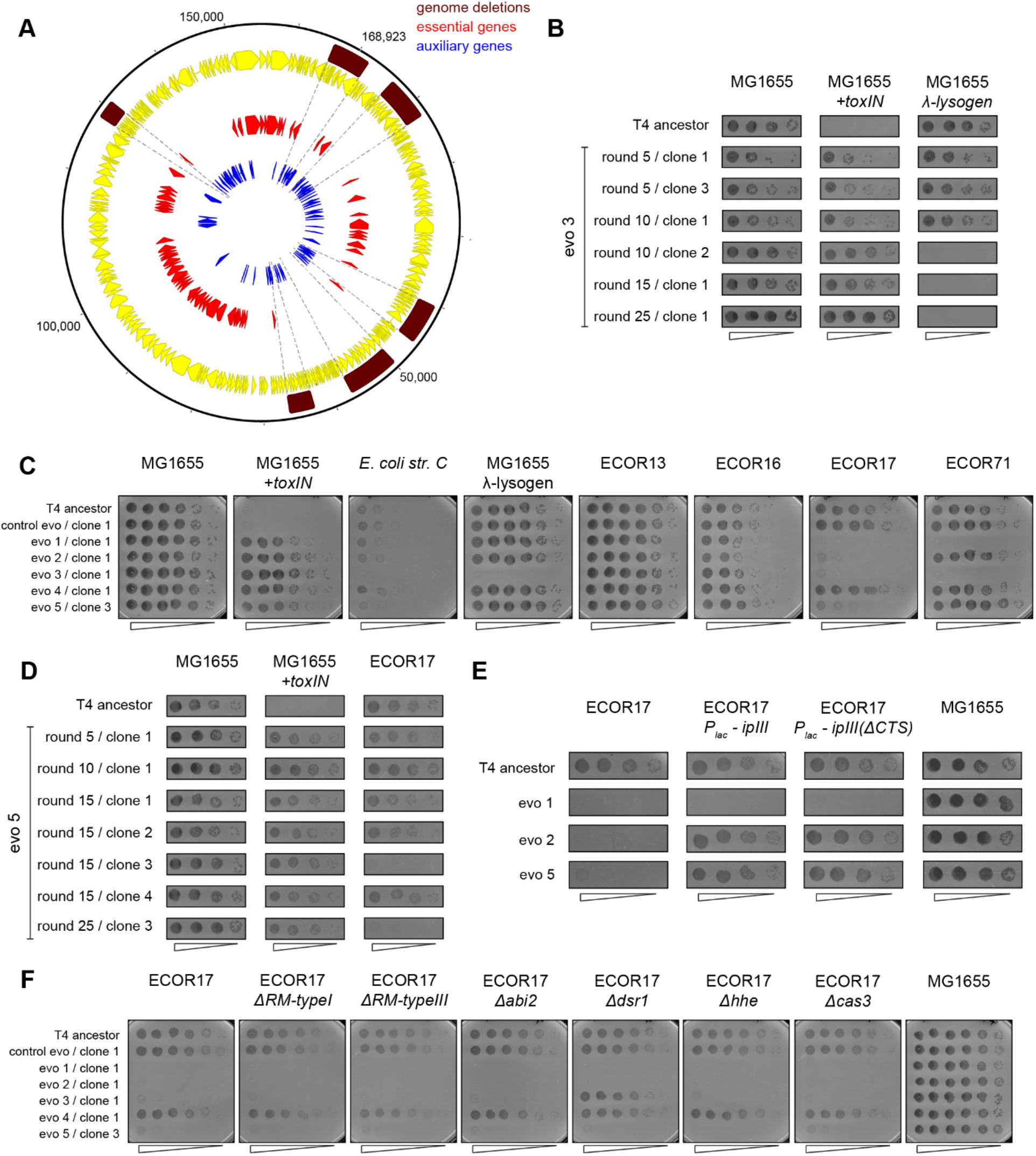
Evolved clones of T4 fix genomic deletions that contain genes essential for infecting alternate hosts. (**A**) Genome map of T4 with all annotated T4 genes (yellow), highlighting essential (red) and auxiliary (blue) genes. Genome deletions identified in evolved populations do not contain any essential genes. (**B**) Serial dilutions of T4 evolved clones isolated from evo 3 after various rounds of evolution (5, 10, 15, 25) spotted on MG1655, *+toxIN*, and λ-lysogen lawns. (**C**) Serial dilutions of T4 evolved clones (from control evo and evo 1 to evo 5) after round 25 and T4 ancestor spotted on *E. coli* strains sensitive to T4 ancestor. These data (performed in triplicate) were quantified to generate the heatmap in Figure 6A. (**D**) Serial dilutions of T4 evolved clones isolated from evo 5 after various rounds of evolution (5, 10, 15, 25) spotted on MG1655, *+toxIN*, and ECOR17 lawns. (**E**) Serial dilutions of T4 evolved clones and ancestor spotted on ECOR17, ECOR17 expressing *IPIII*, ECOR17 expressing *IPIII(ΔCTS)* (mutant that can no longer be packaged into new virions) and MG1655 lawns. (**F**) Serial dilutions of T4 evolved clones (from control evo, evo 1 to 5) after round 25 and T4 ancestor spotted on lawns of ECOR17 with deletions of computationally annotated phage defense systems (*RM type I, RM type III, Abi2, Dsr1, Hhe, CRISPR-Cas type I*).

## References

Anderson, P., and Roth, J. (1981). Spontaneous tandem genetic duplications in Salmonella typhimurium arise by unequal recombination between rRNA (rrn) cistrons. Proceedings of the National Academy of Sciences 78, 3113–3117. https://doi.org/10.1073/pnas.78.5.3113.

Anderson, R.P., and Roth, J.R. (1977). Tandem Genetic Duplications in Phage and Bacteria. Annual Review of Microbiology 31, 473–505. https://doi.org/10.1146/annurev.mi.31.100177.002353.

Andersson, D.I., Slechta, E.S., and Roth, J.R. (1998). Evidence That Gene Amplification Underlies Adaptive Mutability of the Bacterial lac Operon. Science 282, 1133–1135. https://doi.org/10.1126/science.282.5391.1133.

Appelmans, R. (1921). Le dosage du bactériophage. Compt Rend Soc Biol 85, 701.

Bair, C., and Black, L.W. (2007). Exclusion of Glucosyl-Hydroxymethylcytosine DNA Containing Bacteriophages. J Mol Biol 366, 779–789. https://doi.org/10.1016/j.jmb.2006.11.049.

Benzer, S. (1955). Fine Structure of a Genetic Region in Bacteriophage. PNAS 41, 344–354. https://doi.org/10.1073/pnas.41.6.344.

Bernheim, A., and Sorek, R. (2020). The pan-immune system of bacteria: antiviral defence as a community resource. Nat Rev Microbiol 18, 113–119. https://doi.org/10.1038/s41579-019-0278-2.

Blower, T.R., Evans, T.J., Przybilski, R., Fineran, P.C., and Salmond, G.P. (2012). Viral evasion of a bacterial suicide system by RNA-based molecular mimicry enables infectious altruism. PLoS Genet 8, e1003023. https://doi.org/10.1371/journal.pgen.1003023.

Burrowes, B.H., Molineux, I.J., and Fralick, J.A. (2019). Directed in Vitro Evolution of Therapeutic Bacteriophages: The Appelmans Protocol. Viruses 11. https://doi.org/10.3390/v11030241.

Cairns, J., and Foster, P.L. (1991). Adaptive reversion of a frameshift mutation in Escherichia coli. Genetics 128, 695–701. https://doi.org/10.1093/genetics/128.4.695.

Duong, M.M., Carmody, C.M., Ma, Q., Peters, J.E., and Nugen, S.R. (2020). Optimization of T4 phage engineering via CRISPR/Cas9. Scientific Reports 10, 18229. https://doi.org/10.1038/s41598-020-75426-6.

Elde, N.C., Child, S.J., Eickbush, M.T., Kitzman, J.O., Rogers, K.S., Shendure, J., Geballe, A.P., and Malik, H.S. (2012). Poxviruses Deploy Genomic Accordions to Adapt Rapidly against Host Antiviral Defenses. Cell 150, 831–841. https://doi.org/10.1016/j.cell.2012.05.049.

Fineran, P.C., Blower, T.R., Foulds, I.J., Humphreys, D.P., Lilley, K.S., and Salmond, G.P.C. (2009). The phage abortive infection system, ToxIN, functions as a protein-RNA toxin-antitoxin pair. Proc Natl Acad Sci U S A 106, 894–899. https://doi.org/10.1073/pnas.0808832106.

Garb, J., Lopatina, A., Bernheim, A., Zaremba, M., Siksnys, V., Melamed, S., Leavitt, A., Millman, A., Amitai, G., and Sorek, R. (2021). Multiple phage resistance systems inhibit infection via SIR2-dependent NAD+ depletion.

Grossi, G.F., Macchiato, M.F., and Gialanella, G. (1983). Circular Permutation Analysis of Phage T4 DNA by Electron Microscopy. Zeitschrift Für Naturforschung C 38, 294–296. https://doi.org/10.1515/znc-1983-3-422.

Guegler, C.K., and Laub, M.T. (2021). Shutoff of host transcription triggers a toxin-antitoxin system to cleave phage RNA and abort infection. Molecular Cell https://doi.org/10.1016/j.molcel.2021.03.027.

Haldimann, A., and Wanner, B.L. (2001). Conditional-Replication, Integration, Excision, and Retrieval Plasmid-Host Systems for Gene Structure-Function Studies of Bacteria. Journal of Bacteriology 183, 6384–6393. https://doi.org/10.1128/JB.183.21.6384-6393.2001.

Hampton, H.G., Watson, B.N.J., and Fineran, P.C. (2020). The arms race between bacteria and their phage foes. Nature 577, 327–336. https://doi.org/10.1038/s41586-019-1894-8.

Harms, A., Brodersen, D.E., Mitarai, N., and Gerdes, K. (2018). Toxins, Targets, and Triggers: An Overview of Toxin-Antitoxin Biology. Mol Cell 70, 768–784. https://doi.org/10.1016/j.molcel.2018.01.003.

Hendrickson, H., Slechta, E.S., Bergthorsson, U., Andersson, D.I., and Roth, J.R. (2002). Amplification–mutagenesis: Evidence that “directed” adaptive mutation and general hypermutability result from growth with a selected gene amplification. Proc Natl Acad Sci U S A 99, 2164–2169. https://doi.org/10.1073/pnas.032680899.

Hussain, F.A., Dubert, J., Elsherbini, J., Murphy, M., VanInsberghe, D., Arevalo, P., Kauffman, K., Rodino-Janeiro, B.K., Gavin, H., Gomez, A., et al. (2021). Rapid evolutionary turnover of mobile genetic elements drives bacterial resistance to phages. Science 374, 488–492. https://doi.org/10.1126/science.abb1083.

Kauffman, K.M., Chang, W.K., Brown, J.M., Hussain, F.A., Yang, J., Polz, M.F., and Kelly, L. (2022). Resolving the structure of phage–bacteria interactions in the context of natural diversity. Nat Commun 13, 372. https://doi.org/10.1038/s41467-021-27583-z.

Kim, J.-S., and Davidson, N. (1974). Electron microscope heteroduplex study of sequence relations of T2, T4, and T6 bacteriophage DNAs. Virology 57, 93–111. https://doi.org/10.1016/0042-6822(74)90111-1.

Koskella, B., and Brockhurst, M.A. (2014). Bacteria–phage coevolution as a driver of ecological and evolutionary processes in microbial communities. FEMS Microbiology Reviews 38, 916–931. https://doi.org/10.1111/1574-6976.12072.

Kumagai, M., Yamashita, T., Honda, M., and Ikeda, H. (1993). Effects of uvsX, uvsY and DNA topoisomerase on the formation of tandem duplications of the rII gene in bacteriophage T4. Genetics 135, 255–264. https://doi.org/10.1093/genetics/135.2.255.

Kutter, E., Gachechiladze, K., Poglazov, A., Marusich, E., Shneider, M., Aronsson, P., Napuli, A., Porter, D., and Mesyanzhinov, V. (1995). Evolution of T4-related phages. Virus Genes 11, 285– 297. https://doi.org/10.1007/BF01728666.

Leiman, P.G., Kanamaru, S., Mesyanzhinov, V.V., Arisaka, F., and Rossmann, M.G. (2003). Structure and morphogenesis of bacteriophage T4. CMLS, Cell. Mol. Life Sci. 60, 2356–2370. https://doi.org/10.1007/s00018-003-3072-1.

LeRoux, M., Srikant, S., Littlehale, M.H., Teodoro, G., Doron, S., Badiee, M., Leung, A.K.L., Sorek, R., and Laub, M.T. (2021). The DarTG toxin-antitoxin system provides phage defense by ADP-ribosylating viral DNA.

Liebig, H.-D., and Rüger, W. (1989). Bacteriophage T4 early promoter regions: Consensus sequences of promoters and ribosome-binding sites. Journal of Molecular Biology 208, 517–536. https://doi.org/10.1016/0022-2836(89)90145-9.

Mapes, A.C., Trautner, B.W., Liao, K.S., and Ramig, R.F. (2016). Development of expanded host range phage active on biofilms of multi-drug resistant Pseudomonas aeruginosa. Bacteriophage 6, e1096995. https://doi.org/10.1080/21597081.2015.1096995.

Miller, E.S., Kutter, E., Mosig, G., Arisaka, F., Kunisawa, T., and Rüger, W. (2003). Bacteriophage T4 Genome. Microbiol Mol Biol Rev 67, 86–156. https://doi.org/10.1128/MMBR.67.1.86-156.2003.

Mosig, G. (1987). The Essential Role of Recombination in Phage T4 Growth. Annual Review of Genetics 21, 347–371. https://doi.org/10.1146/annurev.ge.21.120187.002023.

Mullaney, J.M., and Black, L.W. (1996). Capsid Targeting Sequence Targets Foreign Proteins into Bacteriophage T4 and Permits Proteolytic Processing. Journal of Molecular Biology 261, 372– 385. https://doi.org/10.1006/jmbi.1996.0470.

Ochman, H., and Selander, R.K. (1984). Standard reference strains of Escherichia coli from natural populations. Journal of Bacteriology 157, 690–693.

Otsuka, Y., and Yonesaki, T. (2012). Dmd of bacteriophage T4 functions as an antitoxin against Escherichia coli LsoA and RnlA toxins. Mol Microbiol 83, 669–681. https://doi.org/10.1111/j.1365-2958.2012.07975.x.

Payne, L.J., Todeschini, T.C., Wu, Y., Perry, B.J., Ronson, C.W., Fineran, P.C., Nobrega, F.L., and Jackson, S.A. (2021). Identification and classification of antiviral defence systems in bacteria and archaea with PADLOC reveals new system types. Nucleic Acids Research https://doi.org/10.1093/nar/gkab883.

Pinilla-Redondo, R., Shehreen, S., Marino, N.D., Fagerlund, R.D., Brown, C.M., Sørensen, S.J., Fineran, P.C., and Bondy-Denomy, J. (2020). Discovery of multiple anti-CRISPRs highlights anti-defense gene clustering in mobile genetic elements. Nat Commun 11, 5652. https://doi.org/10.1038/s41467-020-19415-3.

Pope, W.H., Bowman, C.A., Russell, D.A., Jacobs-Sera, D., Asai, D.J., Cresawn, S.G., Jacobs, W.R., Hendrix, R.W., Lawrence, J.G., Hatfull, G.F., et al. (2015). Whole genome comparison of a large collection of mycobacteriophages reveals a continuum of phage genetic diversity. Elife 4, e06416. https://doi.org/10.7554/eLife.06416.

Rao, V.B., and Black, L.W. (2005). DNA Packaging in Bacteriophage T4. In Viral Genome Packaging Machines: Genetics, Structure, and Mechanism, C.E. Catalano, ed. (Boston, MA: Springer US), pp. 40–58.

Samson, J.E., Magadan, A.H., Sabri, M., and Moineau, S. (2013). Revenge of the phages: defeating bacterial defences. Nat Rev Microbiol 11, 675–687. https://doi.org/10.1038/nrmicro3096.

Stanley, S.Y., and Maxwell, K.L. (2018). Phage-Encoded Anti-CRISPR Defenses. Annual Review of Genetics 52, 445–464. https://doi.org/10.1146/annurev-genet-120417-031321.

Tesson, F., Herve, A., Touchon, M., d’Humieres, C., Cury, J., and Bernheim, A. (2021). Systematic and quantitative view of the antiviral arsenal of prokaryotes. 2021.09.02.458658. https://doi.org/10.1101/2021.09.02.458658.

Wan, H., Otsuka, Y., Gao, Z.Q., Wei, Y., Chen, Z., Masuda, M., Yonesaki, T., Zhang, H., and Dong, Y.H. (2016). Structural insights into the inhibition mechanism of bacterial toxin LsoA by bacteriophage antitoxin Dmd. Mol Microbiol 101, 757–769. https://doi.org/10.1111/mmi.13420.

Wong, S., Alattas, H., and Slavcev, R.A. (2021). A snapshot of the λ T4rII exclusion (Rex) phenotype in Escherichia coli. Curr Genet https://doi.org/10.1007/s00294-021-01183-2.

Wu, D.G., Wu, C.-H., and Black, L.W. (1991). Reiterated gene amplifications at specific short homology sequences in phage T4 produce Hp17 mutants. Journal of Molecular Biology 218, 705– 721. https://doi.org/10.1016/0022-2836(91)90260-D.

Wu, X., Zhu, J., Tao, P., and Rao, V.B. (2021). Bacteriophage T4 Escapes CRISPR Attack by Minihomology Recombination and Repair. MBio 12, e01361–21. https://doi.org/10.1128/mBio.01361-21.

